# Ventral tegmental area dopamine neuron activity mediates multi-valent outcomes during decision making under risk of punishment

**DOI:** 10.64898/2026.07.14.738459

**Authors:** Wonn S. Pyon, Omar A. Viera-Resto, Mojdeh Faraji, Shelby L. Blaes, Caitlin A. Orsini, Max S. Gotlin, Christos Raptis, Carolina Cruz-Wegener, Azin Behnood-Rod, Brandon M. Hellbusch, Sheldon W. Joseph, Jacob Barrett, Hannah M. Holik, Sarthak M. Singhal, Matthew R. Burns, Charles J. Frazier, Jennifer L. Bizon, Barry Setlow

**Affiliations:** Department of Neuroscience, University of Florida, Gainesville, FL, 32610 USA; Department of Psychology, UCLA, Los Angeles, CA, 90095-1564 USA; Department of Psychiatry, University of Florida, Gainesville, FL, 32610 USA; Department of Cellular and Systems Pharmacology, University of Florida, Gainesville, FL, 32610 USA; Department of Neurology, University of Florida, Gainesville, FL, 32610 USA; McKnight Brain Institute, University of Florida, Gainesville, FL, 32610 USA; Center for Addiction Research and Education, University of Florida, Gainesville, FL, 32610 USA

## Abstract

Dopamine contributes to reward-related decision making, but its contributions to decision contexts that include explicit punishment are less well understood. To elucidate the role of ventral tegmental area (VTA) dopamine neurons in decision making under risk of punishment, we used fiber photometry to record activity in these neurons during a decision-making task in which rats choose between a small, “safe” reward and a large reward associated with varying probabilities of explicit punishment. Dopamine neuron activity exhibited phasic increases during risky “Wins” (reward without punishment) and phasic decreases during risky “Losses” (reward plus punishment), each of which scaled with punishment probability and intensity. Further analyses revealed that this outcome-evoked activity predicted choices on subsequent trials. To determine whether VTA dopamine neuron activity plays a causal role in these risk-based decisions, we used optogenetics to selectively inhibit these neurons during risky Wins and Losses. Inhibition of VTA dopamine neurons during Wins selectively reduced the frequency of risky choices after a Win, whereas inhibition during Losses selectively reduced the frequency of risky choices after a Loss. These data reveal that VTA dopamine neuron activity during outcome receipt is causally related to subsequent decision-making behavior. In addition, the fact that the effects of inhibition on choice behavior were specific to the outcome with which the inhibition was paired suggests that VTA dopamine neuron activity contributes to updating after Wins and Losses independently.

**Significance Statement:** Dopamine is implicated in movement, motivation, and learning, and has been strongly linked with substance use and risk-seeking behavior. Despite these associations, it is unclear how the activity of dopamine neurons influences decision making under risk of punishment. To elucidate this, we recorded from ventral tegmental area (VTA) dopamine neurons and found that their activity at the population level integrates reward alongside the probability and intensity of punishment experienced during risky outcomes. Further, inhibition of VTA dopamine neuron activity during risky outcomes decreased risk-seeking behavior in an outcome-specific manner. Our findings align with models suggesting that VTA dopamine neurons signal prediction errors, and provide insight into the role of these neurons in adaptive decision making.

## Introduction

During risk-based decision making, costs are weighed against rewards to guide choices, with the probability, intensity, and nature of the cost being critical moderating variables in the decision-making calculus. Individuals engage in this process on a daily basis and generally make decisions that maximize rewards and minimize costs. Numerous pathological conditions are associated with impairments in this cost/benefit evaluation, however, resulting in maladaptive risk-based decision making. For example, substance use disorder and attention-deficit hyperactivity disorder (ADHD) are characterized by elevated risk-taking behavior in which rewards are overweighted and risks are underweighted. In contrast, individuals with anorexia nervosa display exaggerated risk avoidance due to heightened sensitivity to costs (Bickel et al., 2014; Kaye et al., 2013; Patros et al., 2016). Consequently, a better understanding of the neural mechanisms underlying this form of decision making could be of significant therapeutic benefit.

Dopamine plays a central role in regulating several forms of risk-based decision making, and its contributions within limbic structures including the nucleus accumbens, basolateral amygdala, and prefrontal cortex are well established (Dourado et al., 2025; Freels et al., 2020; Hathaway et al., 2024; Hynes et al., 2020; Mai et al., 2015; Mitchell et al., 2014; Sonntag et al., 2014; St Onge et al., 2011; Truckenbrod, Betzhold, et al., 2023; Wheeler et al., 2024; Zalocusky et al., 2016). In contrast, the contributions to risk-based decision making of ventral tegmental area (VTA) dopamine neurons (which innervate these brain regions) are less well-established. Prior work characterized the function of VTA dopamine activity in decision making involving risk of reward omission, finding that VTA stimulation during a “risky loss” (reward omission after choice of “risky” option) promotes riskier choices, and that inhibition of putative VTA dopamine neurons immediately after a “risky loss” or a “risky win” (reward delivery after choice of “risky” option) shifts choices away from the “risky” option (Stopper et al., 2014). This latter manipulation is ineffective, however, when inhibition occurs after delivery of a “certain” reward (no risk of reward omission). These data suggest that VTA dopamine neurons are necessary to guide choice when evaluating options in terms of both their rewards and their associated risks. The neurobiological mechanisms mediating risk-based decision making can diverge depending on the nature of the risky outcome, however, with distinct substrates governing decision making involving risk of reward omission vs. risk of explicit punishment (Anselme, 2015; Orsini et al., 2015). As such, whether VTA dopamine neurons function similarly during risky decisions when the costs involve explicit punishment remains unknown.

Prevailing theories suggest that VTA dopamine neuron activity signals discrepancies between predicted and actual outcomes (Schultz et al., 1997). Although initially limited to prediction of rewards, other work shows that these neurons signal prediction errors for aversive outcomes (e.g., punishment) with decreases in their activity (Fiorillo et al., 2003; M. Matsumoto & Hikosaka, 2009). More recent work demonstrates that VTA dopamine neurons signal the integrated value of rewarding and aversive outcomes (H. Matsumoto et al., 2016). Dopamine release in NAc, presumably as a consequence of activation of VTA dopamine neurons, also reflects a safety prediction error, with increases in DA occurring when relief from punishment occurs unexpectedly (Stelly et al., 2019). Collectively, these findings provide a strong case for VTA dopamine neurons encoding prediction error-like signals during risk-based decision making in which choice outcomes involve both rewards and probabilistic punishment. To model decision making that incorporates both rewards and risk of punishment in rats, we used the “Risky Decision-making Task” (RDT; Pyon et al., 2025; Simon et al., 2009) in which rats choose between a “safe” option that yields a small reward and a “risky” option that yields a large reward accompanied by varying probabilities of footshock. Task performance is sensitive to punishment probability and intensity, such that selection of the large, risky reward declines with higher probabilities and intensities of footshock (Shimp et al., 2015; Simon et al., 2009). To determine whether VTA dopamine neurons exhibit prediction error-like encoding during this task, we first recorded bulk calcium signals in these neurons in rats during performance in the RDT. We then assessed whether inducing a negative prediction error via optogenetic inhibition of VTA dopamine neurons was sufficient to drive avoidance of risky choices.

## Methods

### Subjects

Male and female (Experiment 1: n=4 male; n=3 female; Experiment 2: halorhodopsin: n=4 male; n=7 female; mCherry: n=4 male; n=3 female) TH-Cre rats on a Long-Evans background (strain: LE-Tg(TH-Cre)3.1Deis strain; RRRC#00659; Witten et al., 2011) were individually housed and maintained on a reversed 12-hour light/dark cycle, with all testing taking place during the dark phase. During behavioral testing, rats were given free access to water, and food restricted to 80-85% of their free feeding weight. Rats were allowed to earn food rewards through task performance and were maintained at their target weights with standard, soy-free laboratory rat chow provided in their home cages. All procedures were conducted in accordance with the Institutional Animal Care and Use Committee of University of Florida and followed NIH guidelines.

### Surgical Procedures

Surgical procedures were performed as in our previous work (Faraji et al., 2025; Hernandez et al., 2019). Rats were given antibiotics (2 mg/kg; subcutaneous) approximately 30 minutes prior to anesthesia and meloxicam (2 mg/kg; subcutaneous) immediately before surgery. Rats were anesthetized with isoflurane gas and secured to a stereotaxic frame. Following a midline incision, the skin was retracted to expose the skull. Four burr holes were then drilled to fit stainless steel anchoring screws for later securing the dental cement (Parkell) and the optic fiber implants to the skull. For photometry experiments, a unilateral burr hole was drilled to microinject 1.0 µL of AAV9-syn-FLEX-jGCaMP8m (∼3.5 × 10^12^ vg/mL titer; AddGene 162378-AAV9; Zhang et al., 2020) into the ventral tegmental area (AP: −5.80, ML: ±2.1 @ 10° medially, DV: −8.7) at a rate of 0.2 µL per minute. After the microinjector was removed and a diffusion period of 5 minutes, a 400 µm diameter optic fiber (NA: 0.66; Doric) was implanted 0.2 mm above the injection coordinates. The left/right position of the optic fiber was counterbalanced across animals. For the optogenetics experiments, bilateral guide cannulae (22-gauge; Plastics One) were implanted targeting the VTA (AP: −5.80, ML: ±2.1 @ 10° medially, DV: −6.7). Microinjectors were then fit to protrude 2.0 mm past the guide cannulae, and 1.0 µL of AAV5-EF1a-DIO-eNpHR3.0-mCherry (3.5 × 10^12^ vg/mL titer; University of North Carolina Vector Core; Gradinaru et al., 2010) was injected into the VTA in each hemisphere at a rate of 0.2 µL per minute. Following removal of the microinjector and a diffusion period of 5 minutes, 200 µm diameter optic fibers (NA: 0.22; ThorLabs) were implanted to protrude 1.0 mm past the end of the guide cannula (DV: −7.7). Following surgery, antibiotics were administered every day for three days, and meloxicam was administered every day for five days.

### Behavioral Apparatus

Both fiber photometry and optogenetic experiments took place in operant chambers housed within sound-attenuating cabinets (Coulbourn Instruments). The cabinets were equipped with 1.12 W houselights located on the interior back walls. The operant chambers each contained a custom food delivery trough (TAMIC Instruments, Gainesville, FL) located in the center of the front wall. The trough projected 3 cm into the chambers and contained a photobeam to detect entries into the trough. A nosepoke port with controllable illumination was positioned 12 cm above each trough and two retractable levers flanking the nosepoke were positioned 11 cm above the floor. The floor of each operant chamber was composed of stainless-steel rods that were attached to scrambled footshock generators (Coulbourn Instruments), and an optical rotary joint was mounted in the ceiling. All operant chambers were interfaced with a computer running Graphic State 4 software that controlled task events and data collection. During fiber photometry, behavioral events such as trial onset, lever presses, and footshock delivery were timestamped by way of TTL pulses sent from the Coulbourn system to the photometry system. During optogenetics experiments, lasers were controlled via TTL pulses sent from the Coulbourn system.

### Experiment 1

#### Behavioral Shaping

After recovery from surgery, rats underwent shaping procedures prior to the RDT and had to meet criterion for each stage of shaping before progressing to the next stage. Rats were initially given 5 of the food pellets used in the behavioral tasks (45 mg soy free pellets, 5TUL, TestDiet) in their home cages prior to the start of shaping procedures to reduce neophobia.

Shaping began with magazine training, in which a houselight illuminated the chamber at the start of the session and a single food pellet was delivered into the food trough every 100 seconds (on average) across a 64-minute session. To progress to the next stage of behavioral shaping, rats had to enter the food trough at least 100 times in a single session. Upon meeting this criterion, rats progressed to lever press training, in which each trial began with extension of one lever (either left or right; counterbalanced between groups) into the operant chamber. Rats received a single food pellet upon pressing the lever. Rats had to make 50 presses in a single 30-minute session to proceed to lever shaping on the opposite lever. After reaching lever pressing criterion on each lever individually, rats proceeded to sessions in which both levers were presented in an alternating fashion, and they were required to make at least 30 presses on each lever within a 30-minute session. Upon reaching criterion, rats moved on to the next shaping stage in which they were trained to nosepoke into the illuminated port above the food trough, which led to extension of one of the two levers. A press on the lever caused the houselight to be extinguished and a food pellet to be delivered. Rats had to perform a minimum of 30 presses on each lever in a 60-minute session (Orsini et al., 2018; Simon et al., 2009, 2011). Upon meeting this criterion, rats repeated this training step while tethered to the rotary joint via an optical patch cord. For the final stage of shaping, rats performed the RDT in the absence of footshock delivery (i.e., reward discrimination). Rats transitioned to the RDT once they displayed a preference for the lever associated with the large reward (greater than 80% presses).

#### Risky Decision-making Task

The version of the RDT used for photometry recordings consisted of three 28-trial blocks over a 35-minute period (8 forced choice trials and 20 free choice trials in each block), with each trial lasting no more than 25 seconds. Each trial began with 5 seconds of houselight illumination, after which the nosepoke light was illuminated. A nosepoke resulted in extension of one (forced-choice trial; 8 per block) or both (free choice trial; 20 per block) levers into the chamber. A trial was considered an omission if rats failed to nosepoke within an 8-second window. A press on the lever designated as the small “safe” lever delivered a small food reward (1 food pellet) whereas a press on the lever designated as the large “risky” lever delivered a large food reward (2 food pellets). The large reward was accompanied, however, by varying probabilities of 1-second footshock delivery. The probability of shock delivery accompanying the large reward increased across consecutive blocks of trials in each session (0, 25, and 75%). Consequently, selection of the risky lever resulted in one of two possible outcomes: 1) receipt of the large reward without footshock (designated a “Win” outcome) or 2) receipt of the large reward accompanied by footshock (designated a “Lose” outcome). On forced-choice trials, the probability of shock following a press on the large “risky” lever was dependent across the 4 trials in each block (e.g., in the 25% block, one and only one of the four large reward lever forced-choice trials was accompanied by shock). In contrast, on free-choice trials, the shock probability on each trial was independent of other trials in that block. Trials were considered an omission if rats failed to press a lever within 5 seconds of extension.

Rats performed 5 consecutive sessions of the task at “Low”, “Medium” and “High” footshock intensities (100 µA, 200 µA and 300-350 µA, respectively), with shock intensities increased weekly (Figure S1B). This design was employed so that dopamine neuron activity could be recorded from each rat across multiple shock intensities.

#### Fiber Photometry

Fiber photometry recordings were performed using Tucker-Davis Technologies (TDT) RZ5P and RZ10X systems. Control of two light-emitting diodes (LEDs; 405 nm and 465 nm; Doric) and recordings of photometry signals and timestamped TTL pulses occurred through Synapse software (TDT). Light was delivered into the implanted optic fiber through a patch cord that was tethered to the implant using a phosphor bronze mating sleeve. Calcium-dependent signal was collected using a 465 nm LED, whereas the 405 nm LED was used as an isosbestic control. Signal was passed through dichroic mirrors and amplified with a Doric photoreceiver.

### Experiment 2

#### Behavioral Shaping

After recovery from surgery, rats underwent shaping procedures in a manner identical to those described in Experiment 1. The one exception was that due to bilateral optic fiber implants, rats experienced a total of three sessions of nosepoke shaping to allow for the gradual introduction of tethered bilateral optic fiber patch cords during behavior.

#### In Vivo Validation of Halorhodopsin Efficacy

Following behavioral shaping, the rats were screened on a discrimination task (adapted from Peng et al., 2021) to validate the efficacy of halorhodopsin *in vivo*. In each of the 56-minutes sessions in this task, rats could select between two levers, each of which delivered 2 food pellets. A press on one of the levers (the “active” lever), however, also resulted in a 5-second delivery of 560 nm laser light (20 mW intensity) to the VTA. Rats were informed of the lever-laser contingencies at the beginning of each behavioral session through a series of forced-choice trials in which only one lever was available. After 8 forced-choice trials (4 trials per lever), rats were given 76 consecutive free-choice trials on which both levers were presented and they were free to choose between them. Once the number of presses on the active lever decreased by at least 35% (or after five consecutive sessions with the same lever-laser contingencies), the contingencies were reversed such that the previously inactive lever now delivered laser light concurrent with food reward and the previously active lever no longer delivered laser light (but still delivered food reward). Rats performed this task through at least one reversal. To progress to the RDT, halorhodopsin-expressing rats had to demonstrate an aversion for the active lever following a contingency reversal. This was defined as a suppression ratio of less than 0.30 calculated as [Post-Reversal Presses/(Pre-Reversal Presses + Post-Reversal Presses)]. This reduction had to occur within five consecutive sessions. Halorhodopsin-expressing TH-Cre rats that failed to meet this criterion were excluded from subsequent testing. A subset of mCherry-expressing TH-Cre rats was tested in the same lever discrimination task using identical procedures, to confirm the lack of a behavioral effect due to laser delivery.

#### Risky Decision-making Task

Rats were trained in a version of the RDT modified for optogenetic manipulations. The task structure was identical to that used in Experiment 1, with a few exceptions. Rather than 25 seconds in duration, each trial was 40 seconds in duration, resulting in a 56-minute session (Orsini et al., 2017). In addition, rats were given 10 seconds to nosepoke or lever press before the trial was scored as an omission. Rats’ choice preference in the RDT exhibits substantial heterogeneity that is stable over time (Mitchell et al. 2014). As such, shock intensity was adjusted individually for each rat, such that they showed roughly 20, 13, and 6 choices of the large reward in the 0, 25, and 75% risk blocks, respectively, as in our previous work (Orsini et al. 2017). Rats were trained in this task until stable performance was achieved (see Data Analysis section for definition of stable performance). They were then assigned to three different optogenetic test conditions, using a within-subjects design such that each rat underwent each test condition in a randomized order. Between each optogenetic session, rats were re-trained in the RDT to re-establish baseline before the next optogenetic session. Optogenetic sessions consisted of tests in which light delivery occurred during either: 1) delivery of the large, unpunished reward (hereafter referred to as “Wins”) 2) delivery of the large, punished reward (hereafter referred to as “Losses”), and 3) delivery of the small, safe reward. Light was delivered for 5 seconds for each condition. Note that not all rats were tested under each of the three conditions due to attrition.

#### Optogenetics

During behavioral testing, laser light (560 nm, 20 mW output at the fiber tip, Shanghai Laser & Optics Century) was delivered bilaterally into the VTA. In order to reach the VTA, light from the laser was passed through a patch cord (200 µm core, Thor Labs) and a rotary joint (1 x 2, 200 µm core, Doric Lenses) mounted on the ceiling of the operant chamber that split into 2 patch cords (200 µm, 0.22 NA, Thor Labs) that connected to the optic fibers implanted in the VTA. The laser was interfaced with the behavioral control hardware/software (Coulbourn Instruments) to allow synchronization of light delivery with task events (Orsini et al., 2017).

#### Ex Vivo Electrophysiology

Whole cell patch clamp electrophysiology was used to verify halorhodopsin function in the VTA of a subset of rats that underwent viral injection surgery in the absence of fiber implantation. At least 4 weeks after surgery, rats were anesthetized using an intraperitoneal injection of 5-10 mg/kg xylazine and 75-100 mg/kg ketamine, and decapitated using a guillotine. The brains were extracted quickly and 300 µm thick coronal sections containing the VTA were made using a Leica VT 1000s vibratome. During sectioning, the brain was submerged in ice-cold sucrose-laden oxygenated aCSF containing the following (in mM): 2 KCl, 1.25 NaH_2_PO_4_, 1 MgSO_4_, 10 D-glucose, 1 CaCl_2_, 206 sucrose, and 25 NaHCO_3_. Slices were then incubated for 30min at 37°C in aCSF which contained the following (in mM): 124 NaCl, 2.5 KCl, 1.23 NaH_2_PO_4_, 3 MgSO_4_, 10 D-glucose, 1 CaCl_2_ and 25 NaHCO_3_. Slices were allowed to equilibrate to room temperature for a minimum of 30 minutes before being used for experiments. The solutions were saturated with 95% O_2_ / 5% CO_2_ to maintain a pH of 7.3. For whole-cell patch-clamp recordings, the slices were transferred to a slice chamber and were continuously perfused at a rate of 2 ml/min with an aCSF bath solution that contained the following (in mM): 126 NaCl, 3 KCl, 1.2 NaH_2_PO_4_, 1.5 MgSO_4_, 11 D-glucose, 2.4 CaCl_2_, and 25 NaHCO_3_. This solution was also saturated with 95% O_2_ / 5% CO_2_ to maintain a pH of 7.3, and bath temperature was maintained at 30-32 ⁰C. Slices were then visualized using infrared differential interference contrast microscopy with an Olympus BX51WI upright stereomicroscope, a 12-bit IRC CCD camera (QICAM Fast 1394, QImaging), and a 40x water-immersion objective. The patch pipettes were prepared with a Flaming/Brown type pipette puller (Sutter Instrument, P-97) from 1.5 mm/0.8 mm borosilicate glass capillaries (Sutter Instrument) and pulled to a tip resistance of 4-7 MΩ. The whole-cell patch-clamp recordings were performed using an Axon Multiclamp 700B amplifier (Molecular Devices). Data were collected at 20 kHz, filtered at 2 kHz, and recorded to a digital computer using a Digidata 1440A and Clampex v10 (Molecular Devices). Tyrosine hydroxylase (TH)-containing neurons in the VTA that expressed mCherry were identified using epifluorescence microscopy (XF102-2 filter set, Omega Optical, excitation 540-580 nm, emission: 615-695 nm). The light source for epifluorescence microscopy was an X-Cite Series 120Q (Lumen Dynamics). Whole-cell patch-clamp recordings were obtained using a potassium-based internal solution that contained the following (in mM): 130 K-gluconate, 10 KCl, 5 NaCl, 2 MgCl_2_, 0.1 EGTA, 2 Na_2_-ATP, 0.3 NaGTP, 10 HEPES, and 10 phosphocreatine, pH adjusted to 7.3 using KOH and volume adjusted to 285-300 mOsm. Halorhodopsin was activated using 1000 ms light pulses, delivered through the excitation filter in the XF102-2 filter set. Experiments were performed in voltage clamp (at −70 mV), in current clamp (at I = 0), or in current clamp during 100-200 pA current injection that was sufficient to drive action potentials. Data were analyzed using custom software written in OriginC (OriginLab, Northampton, MA) by C.J.F. (Orsini et al., 2017).

### Immunohistochemistry

#### Experiments 1 and 2

Rats were injected with 1 mL of Euthasol and then underwent transcardial perfusion with cold 0.1 M PBS followed with cold 4% PFA. Brains were then extracted and postfixed in 4% PFA for 24 hours, then placed in 0.1 M PBS + 20% sucrose solution until cryostat sectioning. Brains were sectioned at 30 µm through the area of the VTA using a cryostat. Coronal sections were collected in a 1-in-4 series and placed in 0.1 M PBS + 0.01% sodium azide-filled wells. Immunohistochemistry was performed on free-floating sections. The sections were incubated in 3% normal donkey serum (Jackson ImmunoResearch Laboratories 017-000-121, 0.3% Triton X-100 and 0.1 M PBS for 1 hour at room temperature on an orbital shaker. Next, the sections were transferred to a primary antibody mixture specific to the antigens of interest (photometry: rabbit anti-GFP (1:1000; Enquire Bioreagents AB29050UL) and sheep anti-TH (1:1000; Invitrogen PA1-4679); optogenetics: rabbit anti-mCherry (1:1000; Invitrogen PA5-34974) and sheep anti-TH (1:1000; Invitrogen PA1-4679)) in 3% normal donkey serum, 0.3% Triton X-100 and 0.1M PBS, and left on the shaker for 48 hours at 4°C. Following the primary antibody, sections underwent three washes in 0.1 M PBS for 10 minutes each after which they were incubated in a secondary antibody mixture (photometry: donkey anti-rabbit conjugated to AlexaFluor-488 (1:500; Invitrogen A201206) and donkey anti-sheep conjugated to AlexaFluor-594 (1:500; Abcam ab150180); optogenetics: donkey anti-rabbit conjugated to AlexaFluor-594 (1:500; Invitrogen A21207) and donkey anti-sheep conjugated to AlexaFluor-488 (1:500; Abcam ab150177)) in 3% normal donkey serum, 0.3% Triton X-100 and 0.1 M TBS) for 2 hours at room temperature on an orbital shaker. Tissue underwent three final 10 minute washes in 0.1 M PBS prior to being mounted on electrostatic slides. Slides were then coverslipped with Prolong Gold Antifade Mountant. Sections were examined under a Keyence BZ-X Series Slide Scanning Microscope to determine the location of optic fibers. Only rats with selective expression of GCaMP, halorhodopsin, or mCherry in TH neurons in the VTA, accompanied by correctly-targeted fiber placements, were included in subsequent data analyses (Figures 1B, 3B and Extended Data Figures 1-1D, 3-1E).

**Figure 1.**
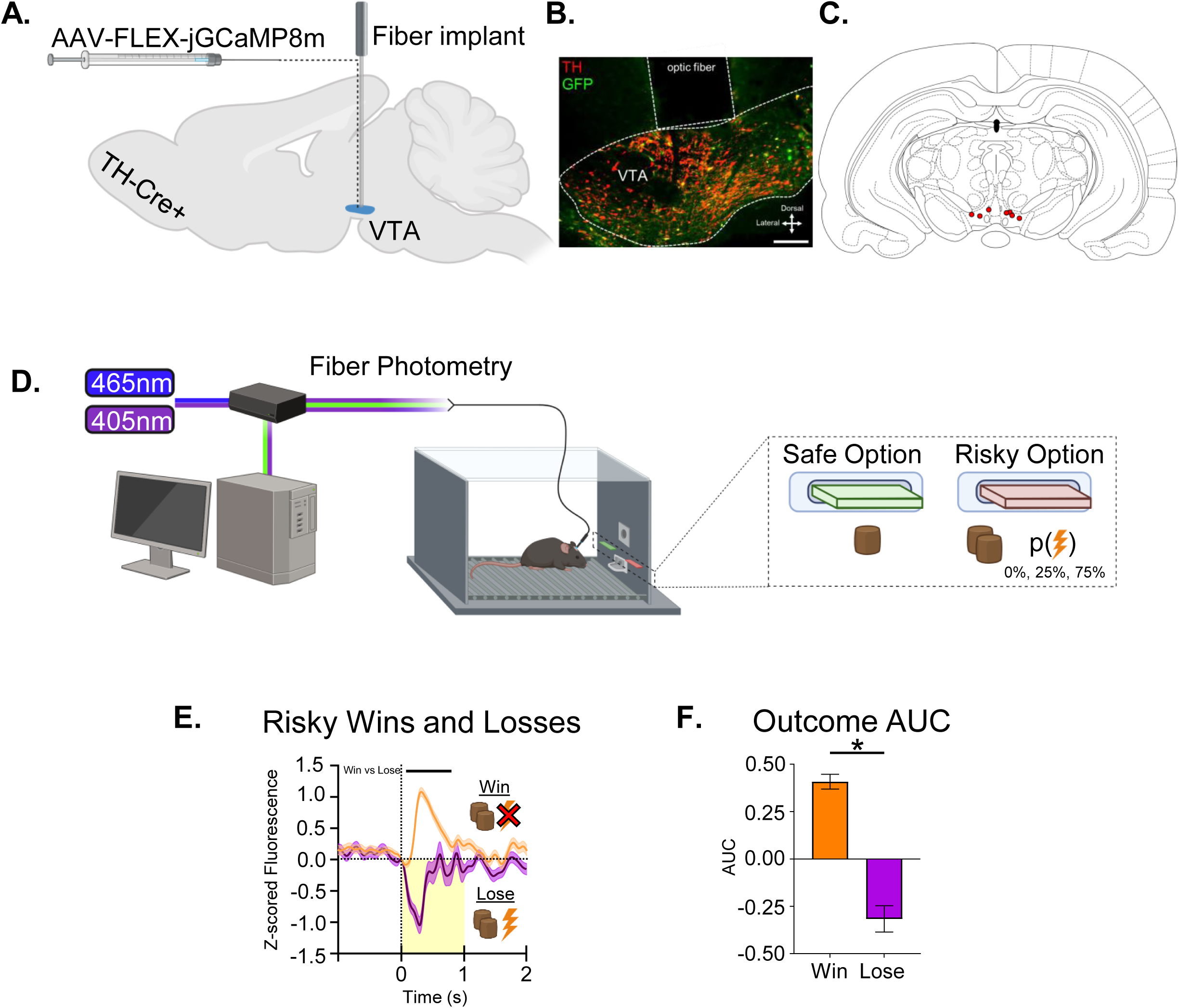
Wins and Losses in the RDT are associated with positive and negative calcium signals, respectively, in dopamine neurons in the ventral tegmental area (VTA). **A)** TH-Cre+ rats were stereotaxically injected with Cre-dependent virus encoding jGCaMP8m into the VTA and implanted with an optic fiber. **B)** Representative image depicting tyrosine hydroxylase (red) and GCaMP-GFP (green) and placement of the optic fiber (dashed line) in the VTA. Scale bar = 100µm. **C)** Estimation of optic fiber placements across animals used for calcium recording. **D)** Illustration of the setup of photometry recordings during RDT performance. **E)** Traces depicting VTA dopamine neuron population activity during risky outcomes on the RDT. Traces represent the average of all Win and Lose trials in blocks 2 and 3 of the RDT performed at High footshock intensity, aligned to the moment of lever press (dashed line, 0s; period of footshock on Lose trials depicted in yellow shading). Bands around traces depict SEM. Black bar above traces indicates time periods within the 0.8-second window following lever press in which the two waveforms are significantly different. **F)** AUC of z-scored fluorescence signal from 0 - 0.8s after lever press. * p < 0.05

### Experimental Design and Statistical Analysis

Photometry signals were processed using MATLAB code modified from Barker et al. (2017). In brief, the isosbestic control signal (405 nm) and calcium-dependent GCaMP signal (465 nm) were collected at 1017 Hz and downsampled to 101 Hz, after which the 405 nm signal was fit to the 465 nm signal using a least-squares polynomial regression. Change in fluorescence was calculated by subtracting the fitted 405 nm signal from the 465 nm signal. The resulting signal was then partitioned into trials according to a houselight (trial onset) TTL delivered from the Coulbourn system. Trial-by-trial signal was z-scored using the mean and standard deviation calculated from a 5-second window prior to the start of each trial (i.e., 5 seconds prior to houselight onset). Z-score normalized signals were then aligned to lever press TTLs, categorized by trial outcome (Safe or risky Win/Loss) and then further categorized into their respective shock probability blocks and shock intensities. Signals from individual trials were averaged across outcomes, and trial-averaged traces were then plotted with bands representing 95% confidence intervals. Bootstrapped confidence intervals (n = 1000) were used to perform waveform analyses based on methodologies described in Jean-Richard-dit-Bressel et al. (2020). A threshold of 17 consecutive significant datapoints was applied to minimize spurious significant findings. AUC values were calculated using a 0.8-second window starting upon lever press (this window was chosen to be the minimum necessary to encompass both increases and decreases in dopamine neuron activity following risky choices). Maximum and minimum z-score in the 0.8-second window following lever press was calculated using the *max* and *min* function, respectively, in MATLAB. Mann-Whitney or Kruskal-Wallis with Dunn’s *post hoc* tests were performed where applicable to determine significant differences between average AUC or minimum/maximum z-score across blocks or shock intensities. A binary logistic regression was used to determine whether AUC during the outcome of Block 3 trials of the RDT at High footshock intensity predicted choice on the next trial. These trials in these sessions were chosen for this analysis as they contained the greatest range of variability in both choice behavior and AUC.

Lever press data from the optogenetics experiments were analyzed using custom Graphic State 4 analysis templates. Graphs were created using GraphPad Prism 7. For the RDT, the number of selections of the risky lever in each block out of 20 trials per block in which rats chose the large, “risky” lever was analyzed. Rats were trained until they displayed stable choice performance on the RDT, which was defined as 2 consecutive sessions with an average coefficient of variation of ≤25% for percent choice of the risky lever. Trial-by-trial analyses were performed to identify the effects of optogenetic manipulations on RDT performance. Win-Stay behavior was calculated by dividing the number of times a rat selected the risky lever after experiencing a risky Win outcome, by the total number of risky Wins. Lose-Shift behavior was determined by dividing the number of times a rat switched to the safe lever or omitted the following trial after experiencing a risky Lose outcome, by the total number of risky Losses (Figure 2M). Paired t-tests were used to evaluate differences in Win-Stay and Lose-Shift behavior as a function of light condition.

**Figure 2.**
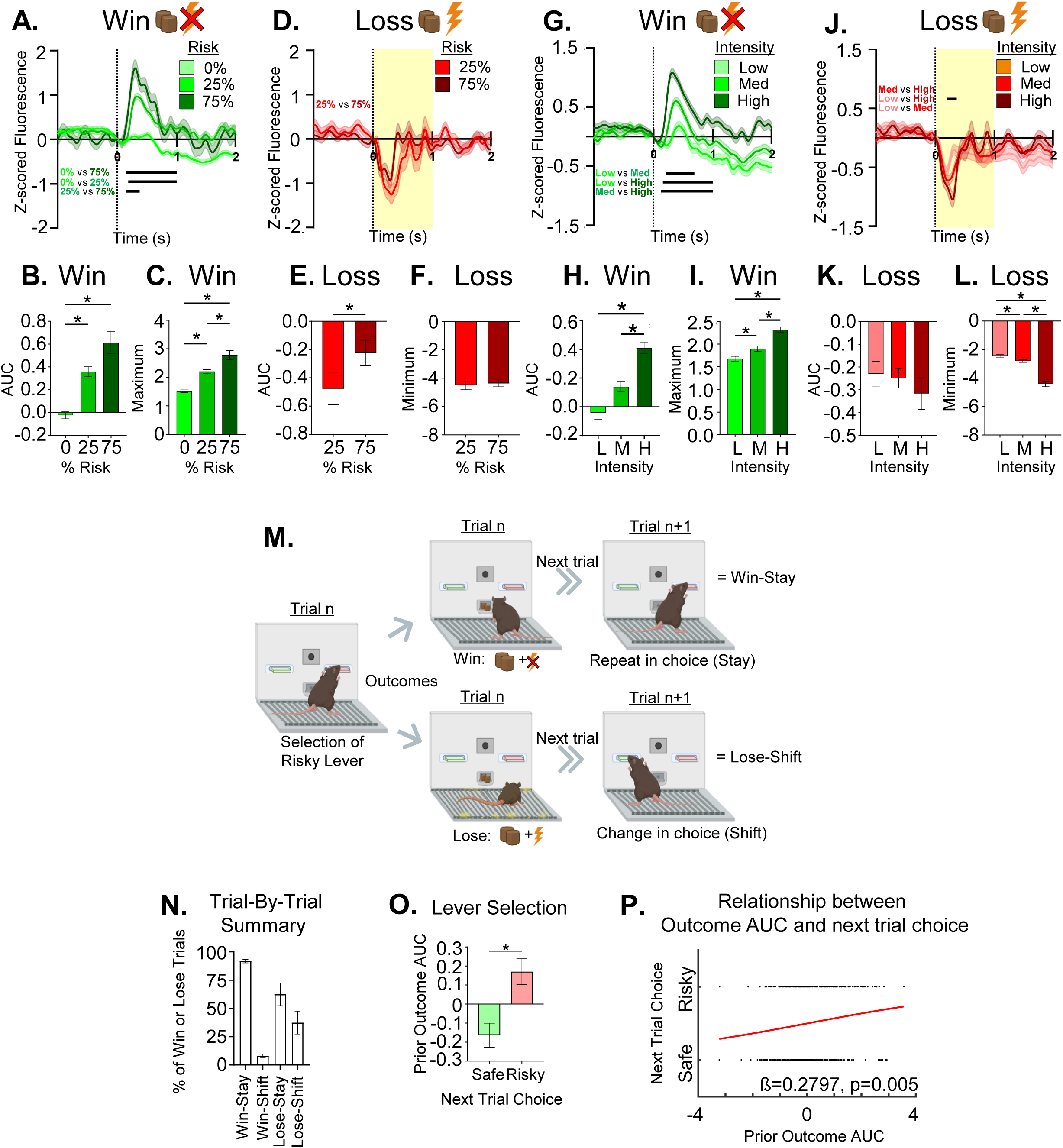
VTA dopamine neuron activity during Win and Lose outcomes in the RDT is modulated by punishment probability and intensity. **A)** Wins experienced at 0%, 25% and 75% risk of High footshock resulted in significantly different VTA dopamine neuron activity. Comparisons of **B)** AUC and **C)** maximum z-score during the 0.8-second window following lever press for Win outcomes experienced at 0%, 25% and 75% risk of footshock. Panels **D, E,** and **F** show the same information for Losses experienced at High footshock. **G)** Wins experienced during 25% and 75% risk blocks of Low, Medium and High footshock sessions result in significant differences in VTA dopamine neuron activity. Comparisons of **H)** AUC and **I)** maximum z-score during the 0.8-second window following lever press for Win outcomes experienced during 25% and 75% risk blocks of Low, Medium and High footshock sessions. Panels **J, K,** and **L** show the same information for Losses experienced during 25% and 75% risk blocks of Low, Medium and High footshock sessions. **M)** Depiction of Win-Stay/Lose-Shift behavior in the RDT. **N)** Summary of actions following Wins and Losses in the RDT for rats in Experiment 1. **O)** Comparison of the AUC at the time of outcome delivery on the trial immediately prior to Safe and Risky choices in the RDT. **P)** Binary logistic regression depicting the relationship between outcome AUC and choice on the next trial. Bands around traces depict SEM. Black lines on panels A, D, G, and J indicate time periods within the 0.8-second window following lever press in which two waveforms are significantly different. * p <0.05

To validate *in vivo* efficacy of halorhodopsin in the discrimination task, a suppression ratio was calculated to quantify the reduction in presses on the laser-paired lever following reversal. The suppression ratio was calculated as [Post-Reversal Presses/(Pre-Reversal Presses + Post-Reversal Presses)]. One-sample t-tests were used to compare the mean suppression ratio within each virus group against a null value of 0.5.

The results of all statistical analyses are reported in Table 1.

**Table 1.**
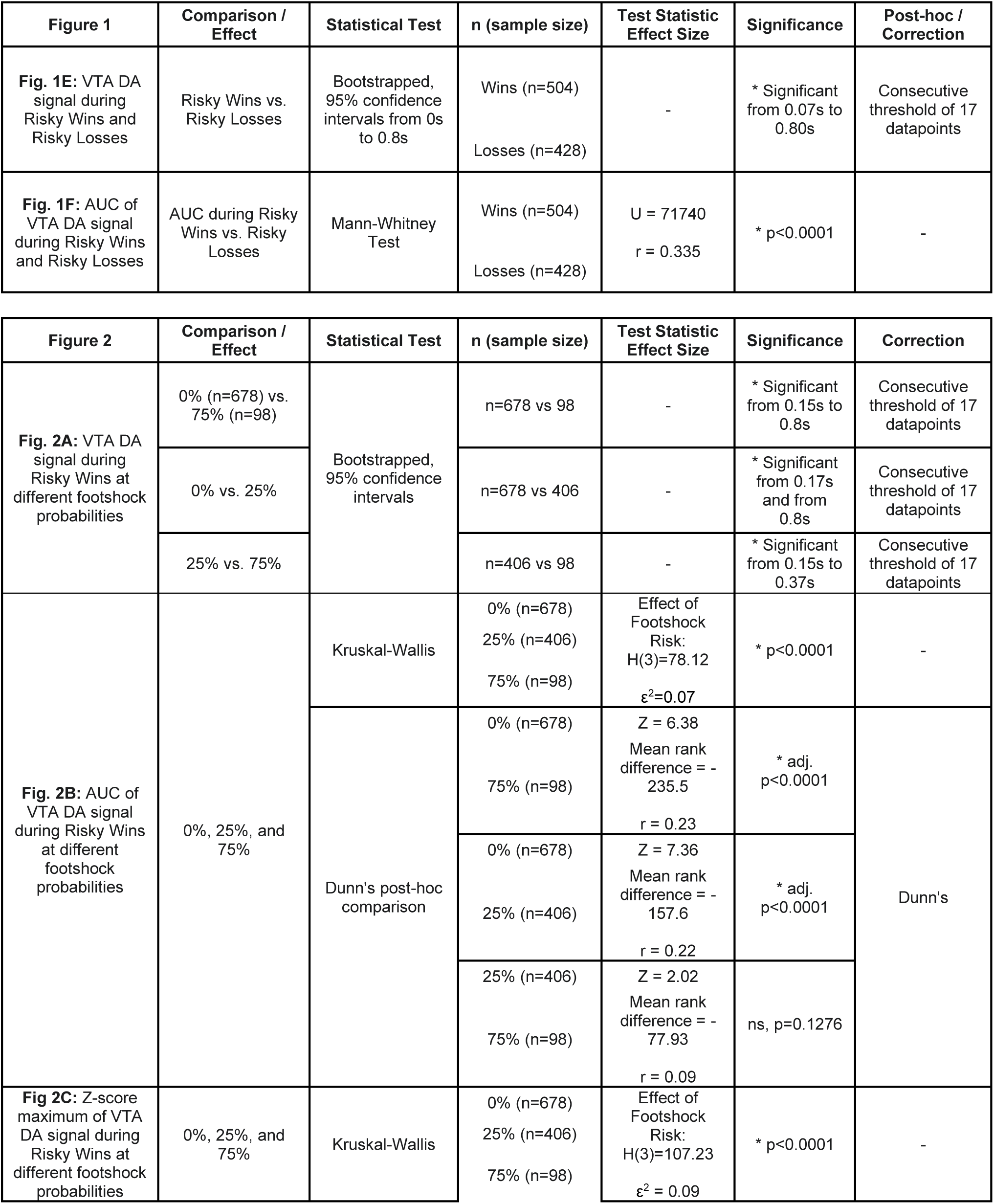
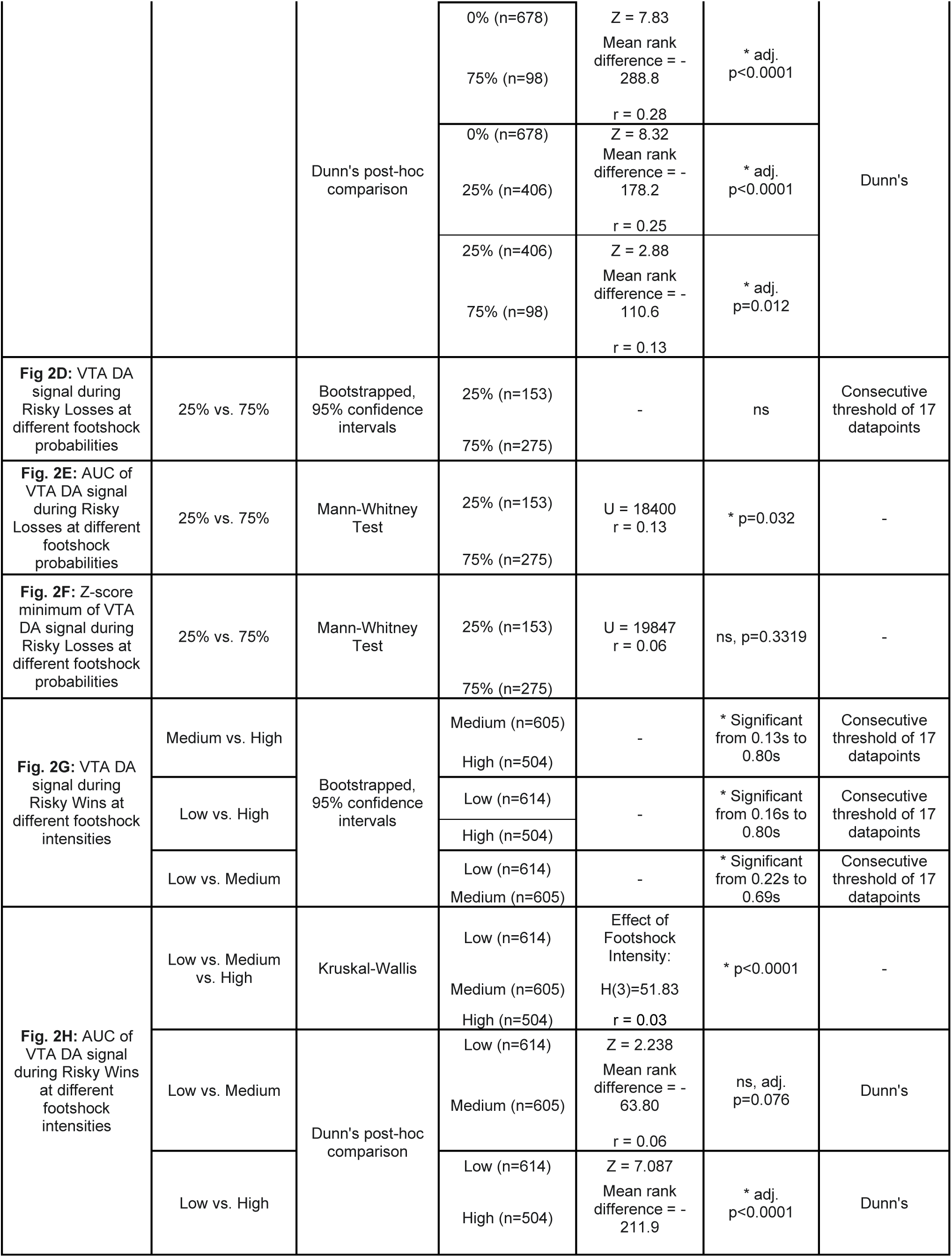
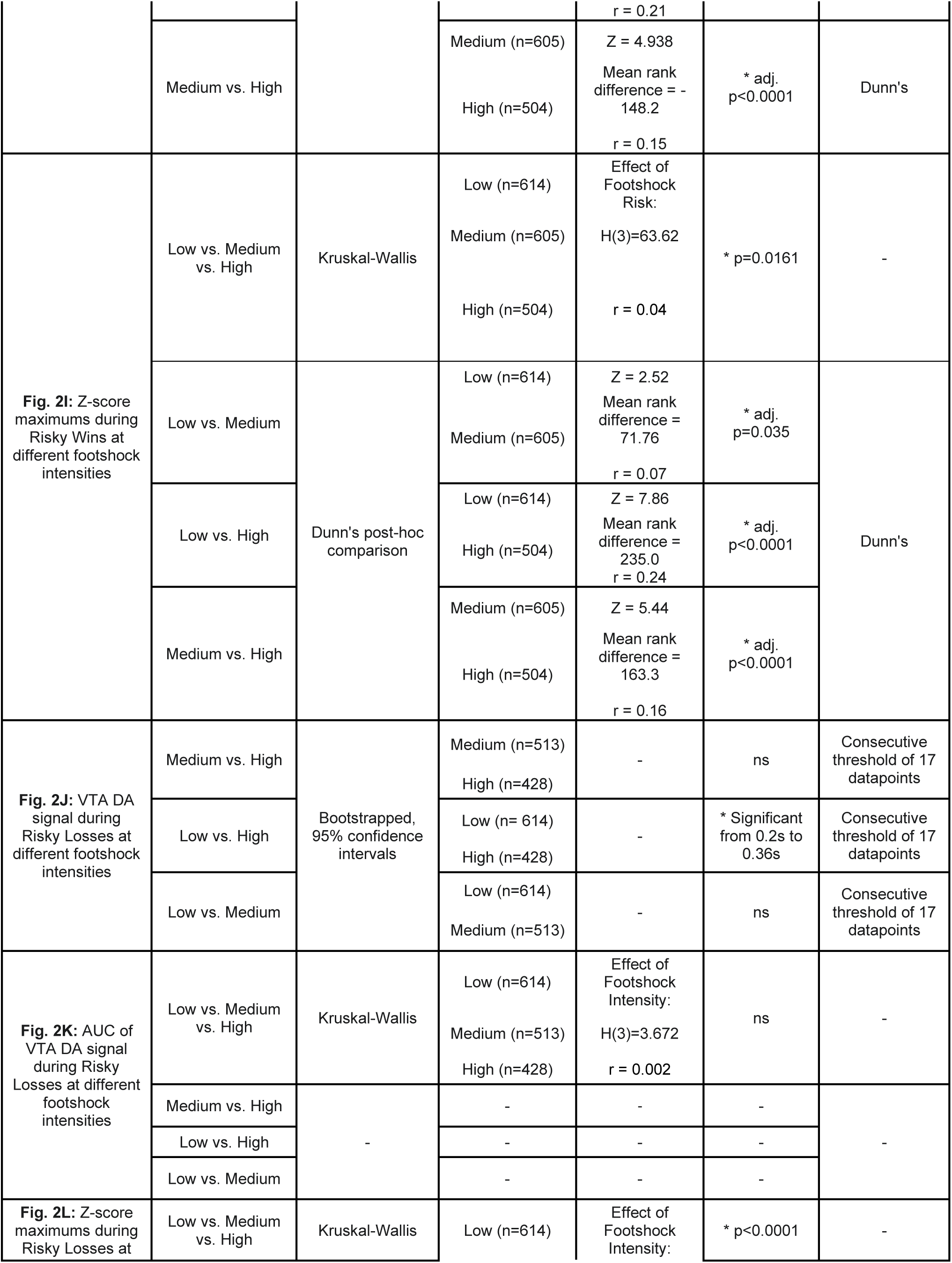
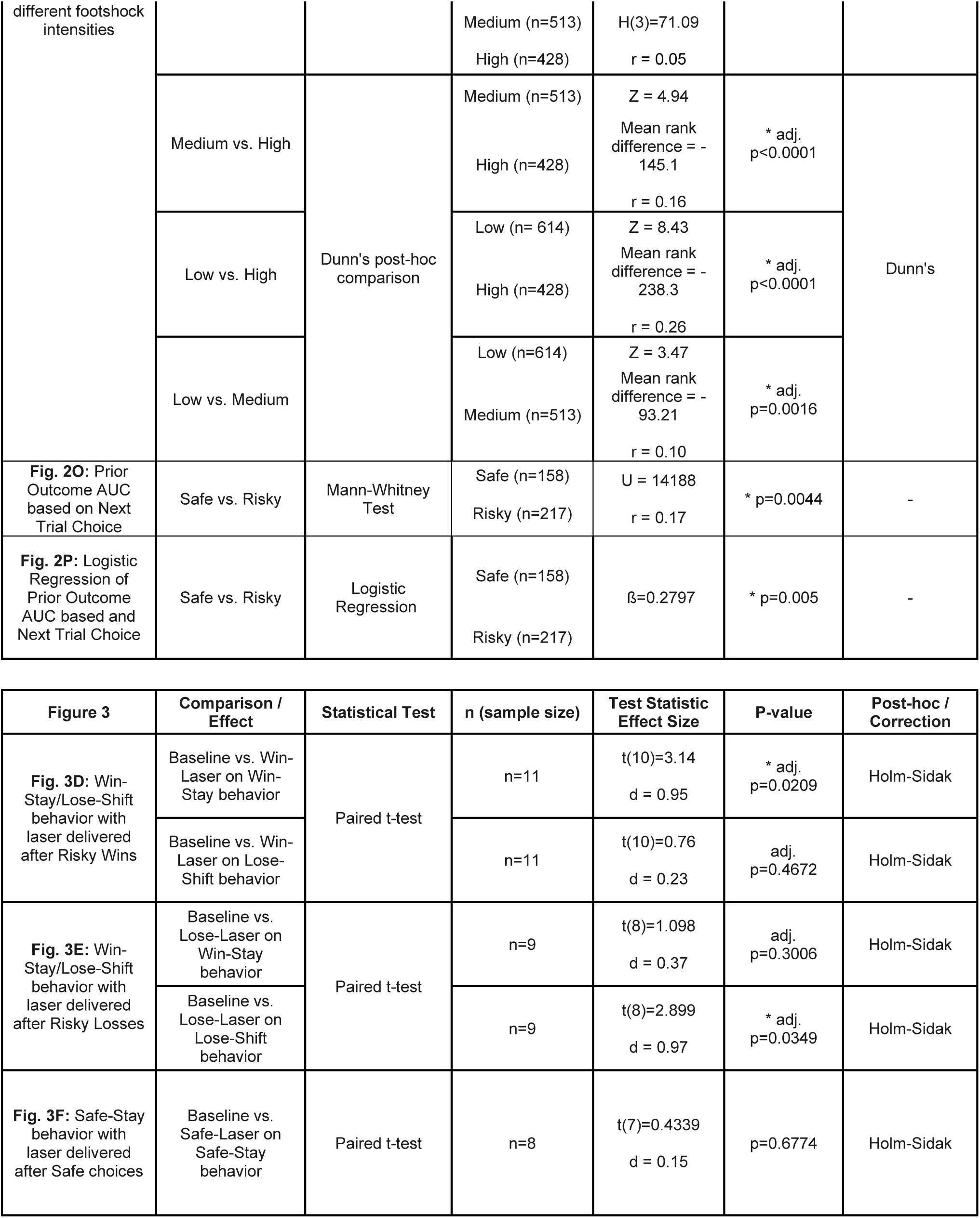
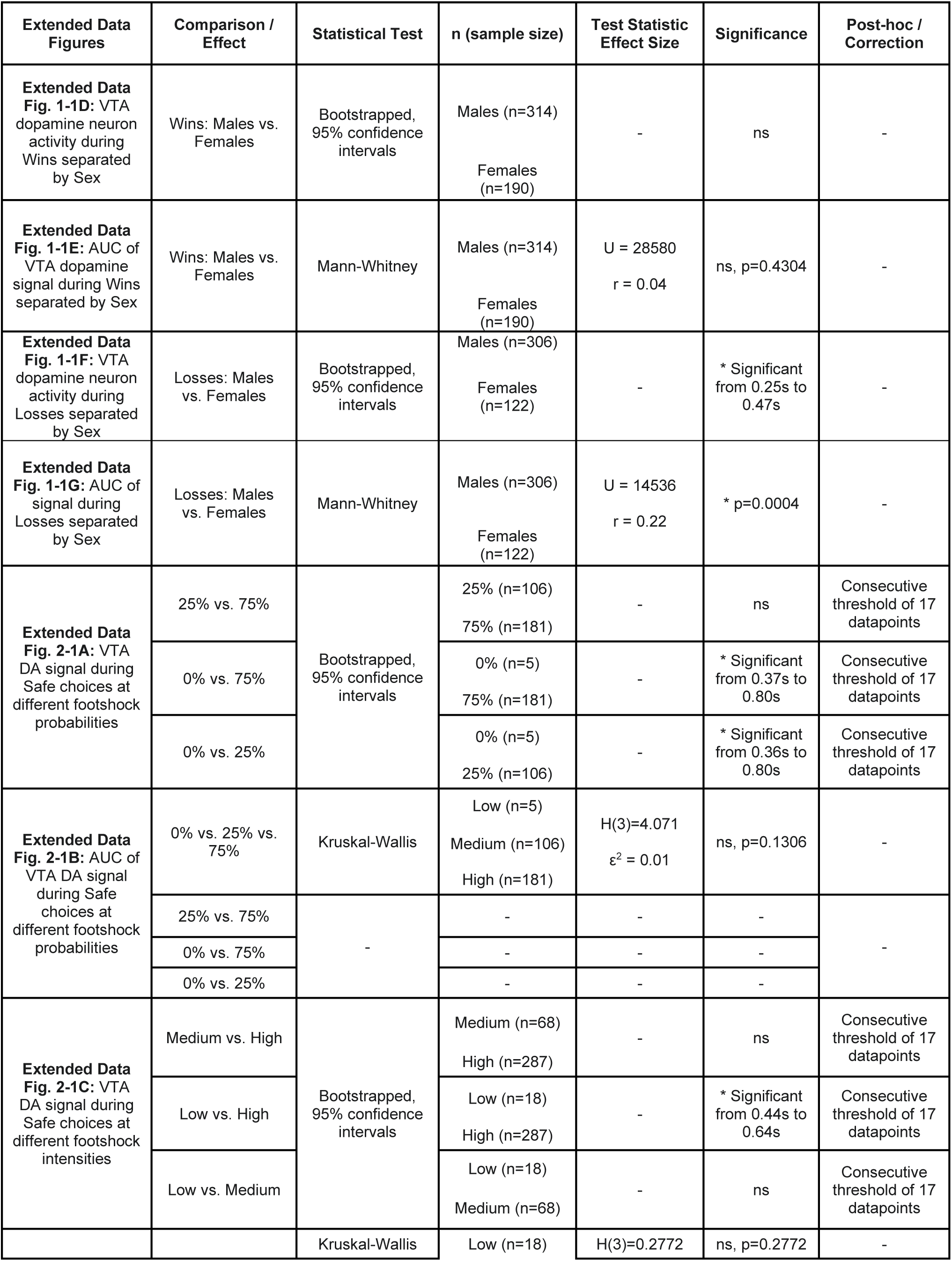
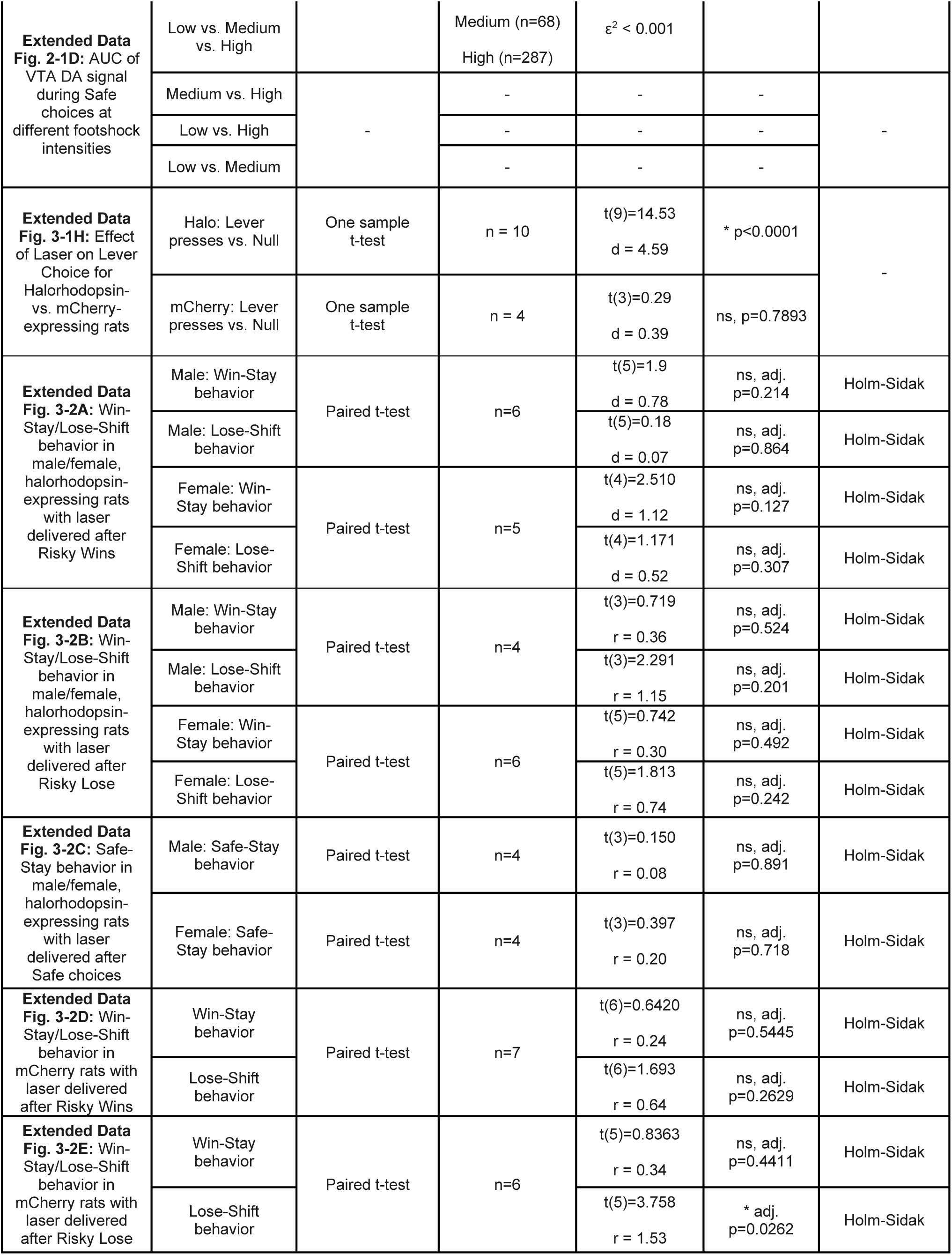
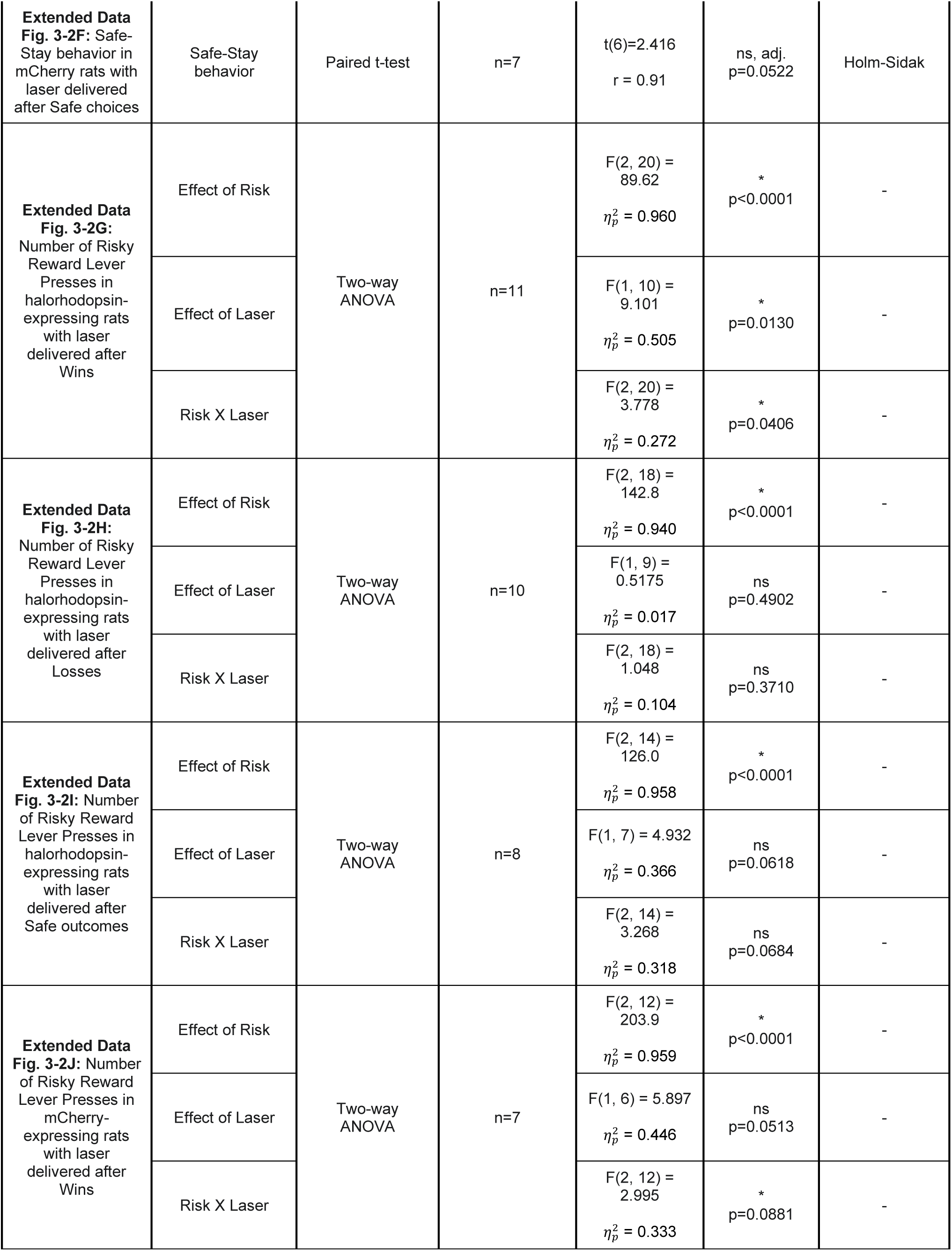
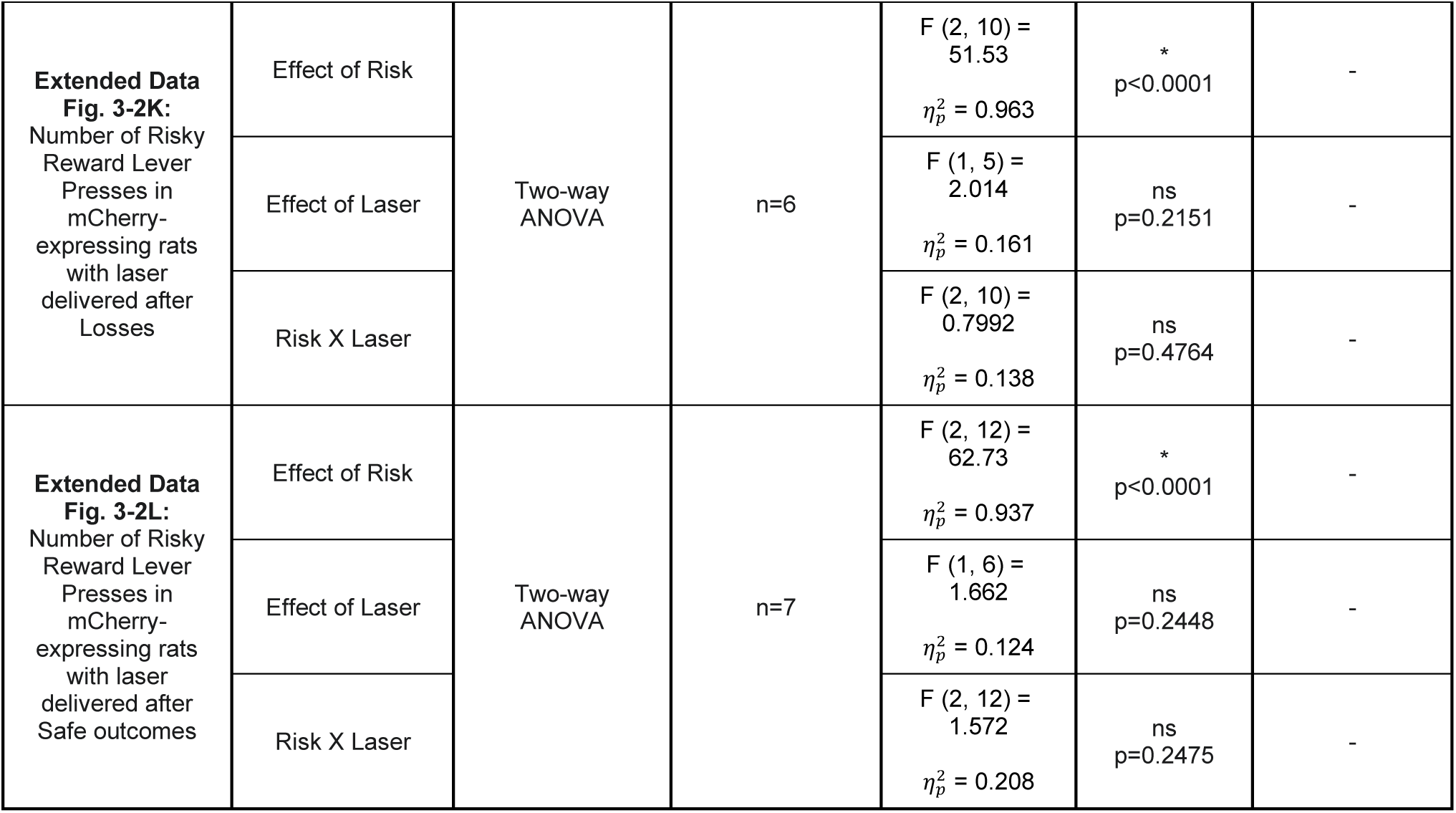
Statistical analyses of experimental results.

## Results

### Experiment 1

#### VTA dopamine neuron population activity exhibits phasic increases and decreases in response to risky Wins and Losses, respectively

To examine the role of VTA dopamine neuron activity and the signaling of prediction errors in risky decision making, calcium activity in VTA dopamine neurons expressing Cre-dependent GCaMP8m (n=7; 4 males, 3 females; Figure 1A-C) was recorded using fiber photometry as rats performed the RDT (Figure 1D). To determine whether VTA dopamine neurons exhibit differential population activity to Win and Lose outcomes, presses on the risky lever in Blocks 2 and 3 spanning at least five days of recording at the High footshock intensity were separated based on outcome (Win or Lose). To account for differences in pre-choice activity that may reflect differences in outcome anticipation across risk blocks, photometry traces were offset such that they crossed the x axis (y=0) at the moment of lever press (x=0). Z-scored fluorescence event-locked to the lever press revealed an increase in VTA dopamine neuron activity after Win trials and a decrease in activity below baseline following Lose trials (Figure 1E). Trial-averaged comparisons of area under the curve (AUC) indicated a significant difference in z-scored fluorescence on Win vs. Lose outcomes (Figure 1F; Mann-Whitney: U = 71740, p <0.0001, r = 0.34). Comparisons of Win and Loss signals between male and female rats revealed no sex differences in the signal and AUC during Wins (Extended Data Figure 1-1E and F and Table 1); however, Losses were associated with a greater decrease in VTA dopamine neuron activity and AUC in females compared with males (Extended Data Figure 1-1G and H and Table 1). Taken together, these results show that VTA dopamine neurons differentially encode positive and negative outcomes involving reward and probabilistic punishment, and that activity in female rats is more sensitive to Losses than in males despite similar activity during Wins.

#### VTA dopamine neuron activity encodes the probability of punishment

The probability of punishment is a key factor in determining whether a benefit is “worth” the risk. It is unclear, however, how the brain encodes information about punishment probability in the context of risky decision making. The RDT is structured such that sessions are segmented into trial blocks that are associated with distinct punishment probabilities organized in an ascending manner. Hence, the probability of Wins (large reward without punishment) decreases while the probability of Losses (large reward with punishment) increases over the session. To determine whether VTA dopamine neuron activity encodes these probabilities, calcium signals on Win and Loss trials following choice of the large, risky option during sessions at High footshock intensity were aligned to lever presses and separated by shock probability block (0, 25, 75%). Trial-averaged traces of z-scored fluorescence underwent waveform analysis (Alonso-Lozares et al., 2024; Taira et al., 2024) to compare z-scored signals across blocks of trials. To reduce spurious significant findings, we required a threshold of at least 17 consecutive significant datapoints (Jean-Richard-dit-Bressel et al., 2020), and limited comparisons to a 0.8-second window following lever press.

Within Win trials, waveform analyses revealed significant differences between z-scored signals across all blocks (Figure 2A and Table 1). Comparisons of AUC (Figure 2B; Kruskal-Wallis test: H(3) = 78.12, p < 0.0001, ε^2^ = 0.07) and z-score maximum (Figure 2C; H(3) 107.2, p < 0.0001, ε^2^ = 0.09) further support these findings and revealed a significant difference in VTA dopamine neuron activity across multiple shock probabilities, with the largest AUC observed on Win outcomes when they are least expected (at the highest probability of punishment, Table 1). While waveform analyses did not reveal significant differences in signal during Losses experienced at the two shock probabilities (Figure 2D), comparisons of AUC for the two signals revealed significant differences, in that the decrease in VTA dopamine neuron activity was significantly larger when animals experienced footshock at low vs. high probability (Figure 2E; Mann-Whitney: U = 18400, p < 0.0001, r = 0.125). Analyses of z-score minimums during this time period did not reveal significant differences (Figure 2F; Mann-Whitney: U = 19847, p = 0.332, r = 0.057). Although VTA dopamine neuron activity differed between safe outcomes experienced in blocks with risk (25% and 75%) and no risk of punishment (0%), there was no significant difference in this activity between the 25% and 75% risk blocks (Extended Data 2-1A, B and Table 1). Collectively, these findings reveal that VTA dopamine neuron population activity during Wins and Losses is not binary (i.e., increasing to better-than-expected outcomes and decreasing to worse-than-expected outcomes) but instead is sensitive to the probability of those outcomes.

#### VTA dopamine neuron activity is sensitive to the magnitude of Wins and Losses

Increasing the intensity of punishment is a common means of discouraging unwanted behaviors, but the mechanisms by which such intensity is encoded during risky decision making are unclear. To evaluate the extent to which VTA dopamine neuron activity encodes the intensity of punishment during Lose outcomes, the footshock intensity was systematically increased in the RDT across weeks of test sessions (“Low”, “Medium”, and “High”; Extended Data Figure 1-1BC). Z-scored fluorescence data from the 25% and 75% shock probability blocks were combined and averaged across five days of testing at each shock intensity.

During risky Wins, VTA dopamine neuron population activity differed significantly across all three intensities (Figure 2G). Similar findings were observed for AUC (Figure 2H; Kruskal-Wallis: H(3) = 51.83, p < 0.0001, ε^2^ = 0.030) and z-score maximums (Figure 2I; Kruskal-Wallis: H(3) = 63.62, p < 0.0001, ε^2^ = 0.037), with increasing AUC and z-score maximum when Wins occurred in the context of a higher footshock intensity (Table 1). For risky Losses, waveform analyses revealed a significant difference in z-scored signal during Losses experienced at Low and High footshock intensity (Figure 2J). Although AUC comparisons did not reveal significant differences across intensities (Figure 2K; Kruskal-Wallis: H(3) = 3.672, p = 0.1595, ε^2^ = 0.002), comparisons of z-score minimums revealed significant differences across all intensities (Figure 2L and Table 1; H(3) = 71.09, p < 0.0001, ε^2^ = 0.046). Finally, during Safe outcomes, waveform analyses revealed brief periods of signal difference between footshock intensities (Extended Data Figure 2-1C and Table 1); however, these differences were not reflected in AUC comparisons (Extended Data Figure 2-1D). Collectively, these results indicate that VTA dopamine neuron activity is sensitive to both the probability and magnitude of Wins and Losses.

#### Relationship between VTA dopamine neuron activity during prior Wins and Losses and subsequent choices

The ability to integrate information about outcomes to influence future decisions is a vital component of optimizing risky decision making. As such, we investigated whether activity observed during the outcomes of risky decisions predicted behavior on the next trial. Trial-by-trial behavior was categorized based on the outcome on the prior trial and the lever chosen on the subsequent trial (Figure 2M). This resulted in four categories of trial-by-trial behavior: Win-stay, Lose-shift, Win-shift, Lose-stay. Of these, Win-Stay and Lose-Shift represented the majority of trial types (Figure 2N). All trials were then separated based on the lever chosen in the second trial, with presses on the safe lever and trial omissions considered “Safe” choices and presses on the risky lever considered “Risky” choices. Comparisons of trial outcome AUC prior to Safe or Risky choices revealed a significant difference (Figure 2O; U = 14188, p = 0.0044, r = 0.172) whereby trials prior to Risky choices had significantly greater AUC compared to trials prior to Safe choices. To investigate if prior outcome AUC predicts Safe or Risky choices on the following trial, we performed a binary logistic regression. The results revealed prior outcome AUC to be a significant predictor of next trial choice (Figure 2P; binary logistic regression: β = 0.2797, p = 0.005), suggesting that VTA dopamine neuron activity during the outcome of the prior trial influences subsequent choice behavior.

### Experiment 2

#### Inhibition of VTA dopamine neurons during risky outcomes results in reduced risk-taking behavior

Although the population activity of VTA dopamine neurons during risky outcomes reveals that these neurons are differentially active during positive and negative outcomes and that this activity predicts future risk-taking behavior, it is unclear if this activity plays a causal role in biasing decision making. To address this, we expressed halorhodopsin (or mCherry) in VTA dopamine neurons in a separate group of TH-Cre rats (as in Orsini et al., 2017; Truckenbrod et al., 2023; Figure 3A-C). Halorhodopsin function was validated *ex vivo* via patch clamp electrophysiology (Extended Data Figure 3-1A-D) and selective expression of halorhodopsin in dopamine (TH+) neurons was confirmed via immunohistochemistry (Extended Data Figure 3-1E). In addition, we confirmed that inhibition of VTA dopamine neurons via halorhodopsin was behaviorally effective using a lever discrimination task (Extended Data Figure 3-1F). Halorhodopsin-expressing rats that had a suppression ratio less than 0.3 (Extended Data Figure 3-1G) were then trained on the RDT.

**Figure 3.**
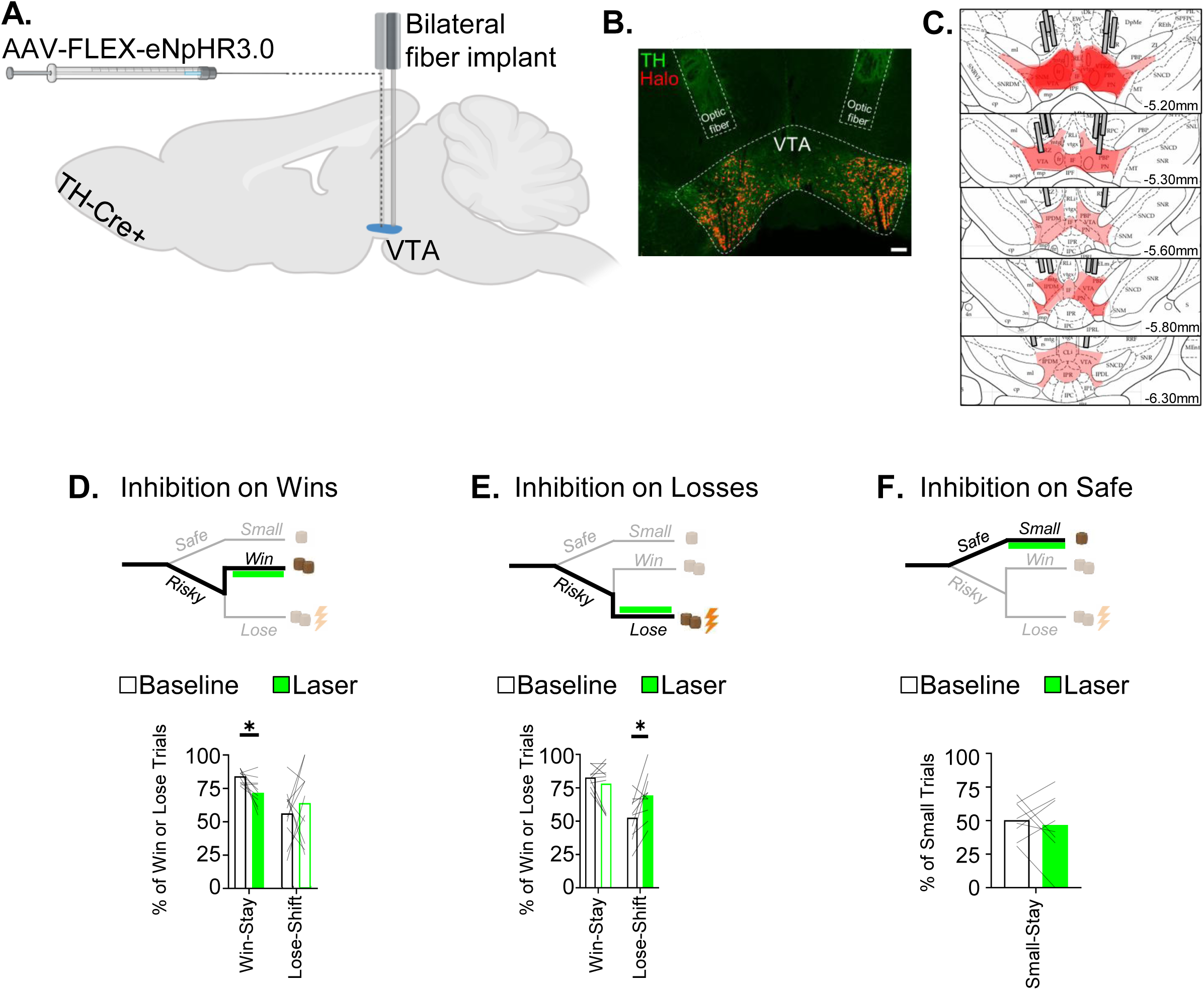
Inhibition of VTA dopamine neurons during Win or Lose outcomes reduces risk-seeking behavior on the next trial. **A)** TH-Cre+ rats were injected bilaterally with Cre-dependent virus encoding halorhodopsin into the VTA and implanted with optic fibers. **B)** Representative image depicting tyrosine hydroxylase (green) and halorhodopsin-mCherry (red) and placement of the optic fibers (dashed lines) in the VTA. Scale bar = 100µm **C)** Estimation of optic fiber placements and halorhodopsin expression across experimental animals. **D)** *Top:* Schematic depicting timing of laser delivery (green bar) during Win outcomes. *Bottom:* Delivery of laser light on Wins significantly reduced Win-Stay behavior. Panels **E** and **F** show the same information for laser delivery on **E)** Losses and **F)** Safe outcomes. * = p < 0.05

After reaching stable baseline performance in the RDT, VTA dopamine neurons were inhibited during delivery of the different outcomes specifically during free-choice trials (Figure 3D-F). Light delivery in halorhodopsin-expressing rats during Wins significantly reduced the proportion of Win-Stay trials compared with baseline (Figure 3D; paired t-test: t(10) = 3.139, p = 0.0209, d = 0.285), with no effect on Lose-Shift trials (paired t-test: t(10) = 0.7558, p = 0.467, d = 0.0687). Light delivery in this condition also significantly reduced the number of choices of the large reward (two-factor ANOVA, main effect of Light Condition: F(1,10) = 9.101, p = 0.013, 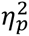 = 0.505; Light Condition x Risk interaction: F(2, 20) = 3.778, p = 0.0406, 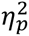 = 0.272; Extended Data Figure S3-2G). In contrast, light delivery during Losses significantly increased the proportion of Lose-Shift trials (Figure 3E; paired t-test: t(9) = 5.758, p = 0.3492, d = 0.576), with no effect on the Win-Stay trials (paired t-test: t(9) 1.098, p = 0.3006, d = 0.1098) and no effect on the number of choices for the large reward (two-factor ANOVA, main effect of Light Condition: F(1,9) = 0.5175, p = 0.4902, 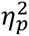 = 0.169; Light Condition x Risk interaction: F(2, 18) = 1.048, p = 0.3710, 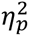 = 0.1044; Extended Data Figure S3-2H). There were no sex differences in the effects of light delivery on any trial type (Extended Data Figure 3-2A-C and Table 1). Finally, light delivery during small, safe outcomes did not affect Safe-Stay behavior (Figure 3F; paired t-test: t(7) = 0.4339, p = 0.6774, d = 0.0542) nor the number of presses for the large reward (two-factor ANOVA, main effect of Light Condition: F(1,7) = 0.4.932, p = 0.0618, 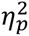 = 0.3658; Light Condition x Risk interaction: F(2, 14) = 3.268, p = 0.0684, 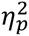 = 0.3183; Extended Data Figure S3-2I).

In mCherry controls rats, there was no effect of light delivery during Wins on either trial type (Extended Data Figure 3-2D). Although light delivery during Losses had no effect on the proportion of Win-Stay trials, it did reduce the proportion of Lose-Shift trials (Extended Data Figure 3-2E). Finally, light delivery during small, safe outcomes did not affect Safe-Stay choice behavior or the number of large reward choices in these rats (Extended Data Figure 3-2F, G and Table 1).

Consistent with our finding that reduced VTA dopamine neuron activity predicts reduced risk taking, the results of this experiment revealed that inhibition of VTA dopamine neuron activity during risky outcomes reduced the frequency of risk-taking behavior on the next trial.

## Discussion

We examined the contributions of VTA dopamine neurons to decision making involving risk of explicit punishment. These neurons increased their activity during risky “Win” outcomes and decreased their activity (below baseline) during risky “Loss” outcomes. Similar to rats’ choice performance, neural activity was sensitive to both the probability and intensity of punishment. These findings are consistent with evidence showing that midbrain dopamine neurons signal prediction errors (Schultz et al., 1997) as well as with subsequent extensions of these findings related to probabilistic food reward (Fiorillo et al., 2003) and airpuffs (H. Matsumoto et al., 2016; M. Matsumoto & Hikosaka, 2009). To determine whether these prediction error-like signals play a causal role in guiding choice behavior, we optogenetically inhibited VTA dopamine neurons during each of the possible choice outcomes (i.e., mimicking the decrease in activity observed following risky “Loss” outcomes). Temporally-specific inhibition of VTA dopamine neurons during risky outcomes (both Wins and Losses) resulted in a shift away from risky choices on subsequent trials. These findings are consistent with prior work showing that VTA neuron inhibition during reward receipt associated with risk of omission attenuates preference for these rewards (Stopper et al., 2014), and extend it to the context of punishment.

Importantly, the effects of VTA dopamine neuron inactivation in the current experiment were specific to the outcome with which it was paired. Inactivation during Losses increased the probability of shifting choices to the safe option following Losses, without a concomitant shift to the safe option following Wins. Similarly, inactivation during Wins decreased the probability of continuing to choose the risky option following Wins, without a concomitant shift in choices following Losses. This dissociation is notable because some theories suggest that devaluation of an outcome would prompt a reduction in the behavior associated with that outcome (Adams & Dickinson, 1981; Balleine & Dickinson, 1998), such that devaluation of either Wins or Losses would result in a global decrease in choice of the risky lever. In contrast, the current results imply that the optogenetically-induced reduction in outcome value was selective to specific conjunctions of outcomes of the risky choice (i.e., the large reward either with or without punishment), rather than a global outcome signal following the risky choice (although note that inactivation during Wins did produce a reduction in choices of the large reward when there was no risk of punishment, perhaps because of the large number of Win trials on which inactivation occurred during this block; Extended Data Figure S3-2G). Notably an outcome-specific effect of optogenetic inactivation in this study corroborates recent findings that VTA dopamine release in the basolateral amygdala supports the encoding of identity-specific reward memories (Sias et al., 2024). Wins and Losses may represent distinct outcome identities associated with the risky lever, and their individual pairing with optogenetic inactivation may devalue one outcome without affecting the other.

An unexpected finding in this work was the fact that light delivery in mCherry-expressing control rats during Lose outcomes promoted more risky choices on the following trial (Extended Data Figure 3-2C). Because no opsin was present in the brains of these animals, this effect was likely the result of light-induced local tissue heating (Christie et al., 2013) whereby even low intensity light delivered through an optic fiber can prompt small but physiologically-relevant increases in temperature and neural excitability (Guatteo et al., 2005; Shapiro et al., 2012). Such heat-induced increases in dopamine neuron activity would be expected to oppose the activity-suppressing effects observed in halorhodopsin animals and perhaps produce reinforcing effects on their own. Effects on behavior in mCherry control animals were not observed with light delivery on risky wins, however, nor were they evident in the *in vivo* optogenetic validation experiment (Extended Data Figures 3-1G and 3-2D), suggesting that light delivery effects in mCherry control rats are relatively subtle. Most importantly, however, the effect of light delivery in mCherry control animals during Lose outcomes in the RDT was in the opposite direction of that in halorhodopsin animals (Figure 3E), suggesting that activation of halorhodopsin was not only effective but sufficient to counteract any effects of light-induced tissue heating.

The fact that VTA dopamine neuron activity responded to unexpected Wins and Losses via increases and decreases in activity that scaled with both probability and intensity of footshock punishment appears to contrast with recent work showing that in Pavlovian conditioning contexts, VTA dopamine neuron activity exhibited only limited increases in response to both cued and uncued footshock (Stelzner et al., 2025). The reasons for this discrepancy are not clear but could include factors such as subtle differences in the specific VTA subregion targeted, the probabilistic vs. deterministic nature of the shock, and/or differential engagement of dopamine neurons in instrumental vs. Pavlovian contexts. It has also been suggested that (at least in the nucleus accumbens) decreases in dopamine signals in response to negative outcomes emerge more slowly during training than increases in dopamine signals in response to positive outcomes (Burke et al., 2026). As such, the fact that rats in the current study were well trained in the RDT could account for the presence of negative prediction error-like signals. Interestingly, the activity of VTA GABAergic neurons recorded in the Stelzner et al. study showed more robust contextual modulation compared to dopamine neurons. Evaluation of the contributions of these GABAergic neurons to decision making under risk of explicit punishment will be an important avenue for future work.

The experiments presented here were not powered to investigate sex differences; nevertheless, a significant sex difference was detected in the activity of VTA dopamine neurons, with females exhibiting a greater change in activity on Loss but not Win trials compared to males. This larger Loss signal in female rats is consistent with their reduced preference for the large, risky reward in the RDT compared to males (Orsini et al., 2016, 2021; Truckenbrod, Cooper, et al., 2023), as well as higher levels of risk aversion and sensitivity to probabilistic punishment more broadly (Chowdhury et al., 2019; Truckenbrod et al., 2025). There were, however, no sex differences in the effects of inactivation of these neurons on choice behavior in Experiment 2, suggesting that the quantitative difference in the magnitude of the decreased VTA dopamine neuron activity in response to Losses does not reflect a qualitative sex difference in the function of this activity change in guiding choice behavior.

Collectively, these experiments show that VTA dopamine neurons play causal roles in decision making under risk of explicit punishment, and are consistent with recent work indicating that decreases in the activity of these neurons are necessary for learning from punishment (Tan et al., 2026). The population-level activity of these neurons signals information concerning multi-valent outcomes in a manner that conveys punishment probability and intensity, both when punishment is delivered and when it is omitted. This activity is then capable of driving trial- and outcome-specific changes in risk-taking behavior, likely in part through signaling to downstream brain regions key for updating task-related relationships and facilitating adaptive behavior (Piantadosi et al., 2026). These findings expand on the limited existing literature investigating the role of VTA dopamine neurons in risky decision making and provide novel evidence of their role in signaling a comprehensive prediction error signal in the context of explicit punishment. It is important to note, however, that the bulk dopamine neuron activity assessed in the present study does not capture the heterogeneity in the response properties of these neurons that is evident in their terminal regions (e.g., Mohebi et al., 2019, 2024). Future research focused on specific dopaminergic populations projecting to distinct limbic-striatal targets may reveal other roles for dopamine in decision making under risk of punishment. In addition, it is unclear whether VTA dopamine neurons engage in novel computations, or whether predictive coding is communicated to the VTA via its afferents (Bouret & Sara, 2004; Feng et al., 2024; Jordan, 2024). Key work highlights upstream brain regions such as the hypothalamus (Barker et al., 2023; de Jong et al., 2019), and habenula (Li et al., 2021; Stopper et al., 2014) as being critical for relaying information about aversive outcomes and associated stimuli to VTA dopamine neurons. How these VTA afferent regions encode aversive experiences that are then utilized for an error computation, however, is unknown.

## Acknowledgements

This research was supported by NIH R01DA036534 (BS), RF1AG060778 (JLB, BS, CJF), K99DA041493 (CAO), T32AG061892 (WSP), and T32DA024635 (WSP).

## Extended Data

**Figure 1-1.**
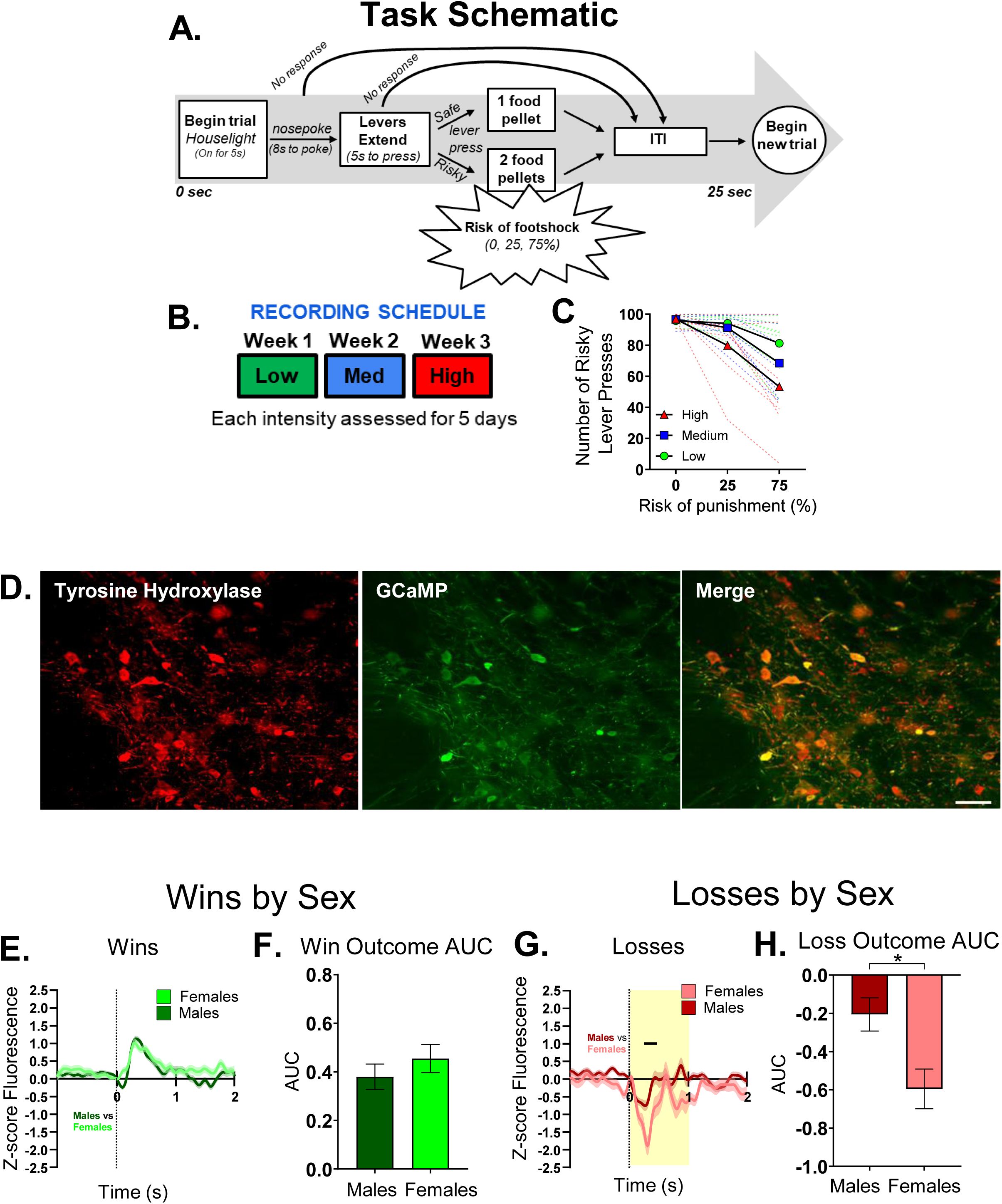
Schematic of the Risky Decision-making Task (RDT), recording schedule, and sex differences in Win/Lose signal in Experiment 1. **A)** Schematic depicting task-relevant details of the RDT. **B)** Rats performed the RDT beginning at Low footshock intensity, ascending to Medium, then High after 5 consecutive sessions at each intensity. **C)** Performance in the RDT (number of choices of the large reward at each risk of punishment) at Low, Medium and High footshock intensities, separated into group mean (bold line) and individual (dashed) values. **D)** Representative image depicting colocalization of tyrosine hydroxylase (red) with GCaMP-GFP (green) in the VTA. Scale bar = 40µm **E)** VTA dopamine neuron calcium signals and **F)** AUC in males and females during Win outcomes. Panels **G** and **H** depict the same information but for Loss outcomes. Bands around traces depict SEM. Black line on panel G indicates time periods within the 0.8-second window following lever press in which two waveforms are significantly different. * p <0.05

**Figure 2-1.**
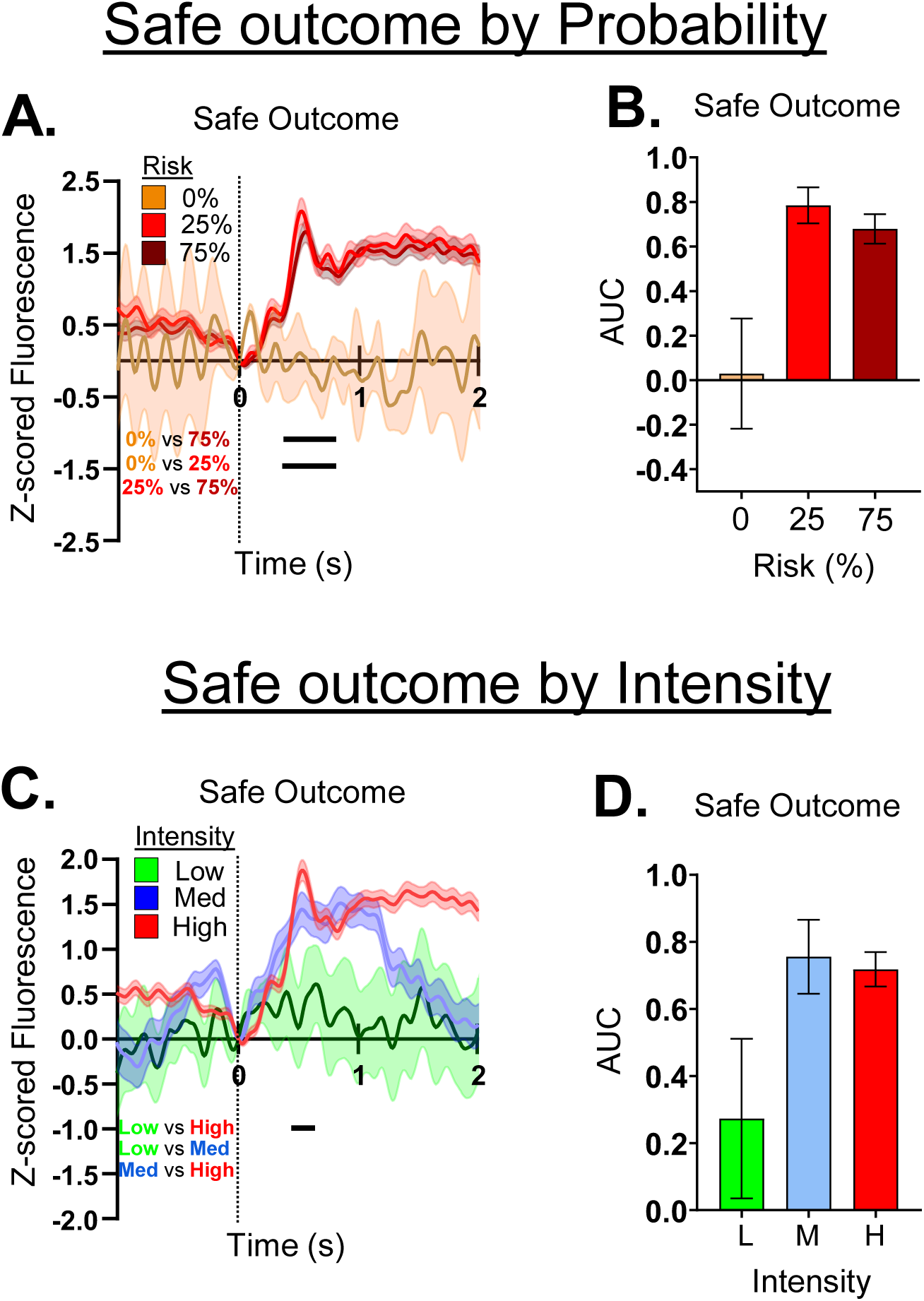
VTA dopamine neuron activity during safe outcomes. **A)** VTA dopamine neuron calcium signals during Safe outcomes following selection of the small reward lever at 0%, 25%, and 75% risk blocks. **B)** AUC of the 0.8-second window following lever press for Safe outcomes during 0%, 25% and 75% risk blocks. Panels **C** and **D** depict the same information but for Safe outcomes experienced when footshock intensity was Low, Medium, and High. Bands around traces depict SEM. Black lines on panels A and C indicate time periods within the 0.8-second window following lever press in which two waveforms are significantly different.

**Figure 3-1.**
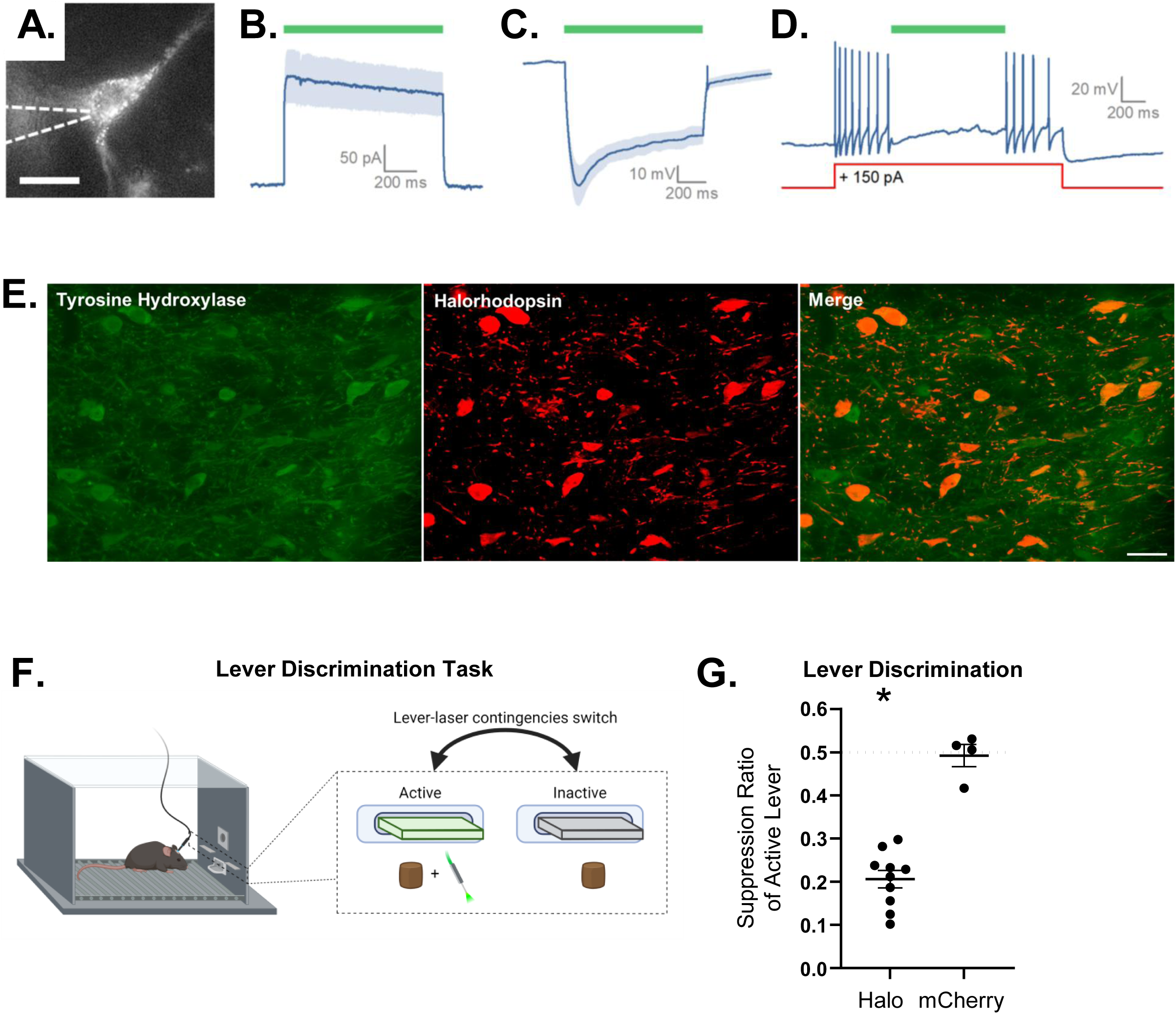
*Ex vivo* and *in vivo* validation of halorhodopsin expressed in dopamine neurons in the VTA. **A)** Halorhodopsin-expressing (mCherry-positive) VTA neurons were identified for whole-cell patch clamp recording using epifluorescence microscopy. Scale bar = 20 µm**. B)** One second exposure to 560nm light produced an outward current in VTA neurons voltage clamped at −70mV (175.12 ± 42.79 pA, n = 5, p=.02). **C)** Identical stimulation produced a hyperpolarization from rest in current clamped VTA neurons (−44.54 ± 5.74 mV, n=5, p=.001). **D)** A representative trace showing that halorhodopsin-mediated hyperpolarization is sufficient to stop action potential firing in current clamp as produced by a 150 pA depolarizing current pulse. **E)** Representative image depicting colocalization of tyrosine hydroxylase (green) with halorhodopsin-mCherry (red) in the VTA. Scale bar = 40 µm **F)** Schematic of the lever-laser discrimination task used to confirm functional efficacy of halorhodopsin *in vivo*. **G)** Quantification of lever-laser discrimination task performance using a suppression ratio. The dashed horizontal line at 0.5 represents the null ratio if laser delivery contingent with lever press had no behavioral effect. Halorhodopsin-expressing animals exhibited significant fewer presses for the active lever compared to the null ratio. * p < 0.05.

**Figure 3-2.**
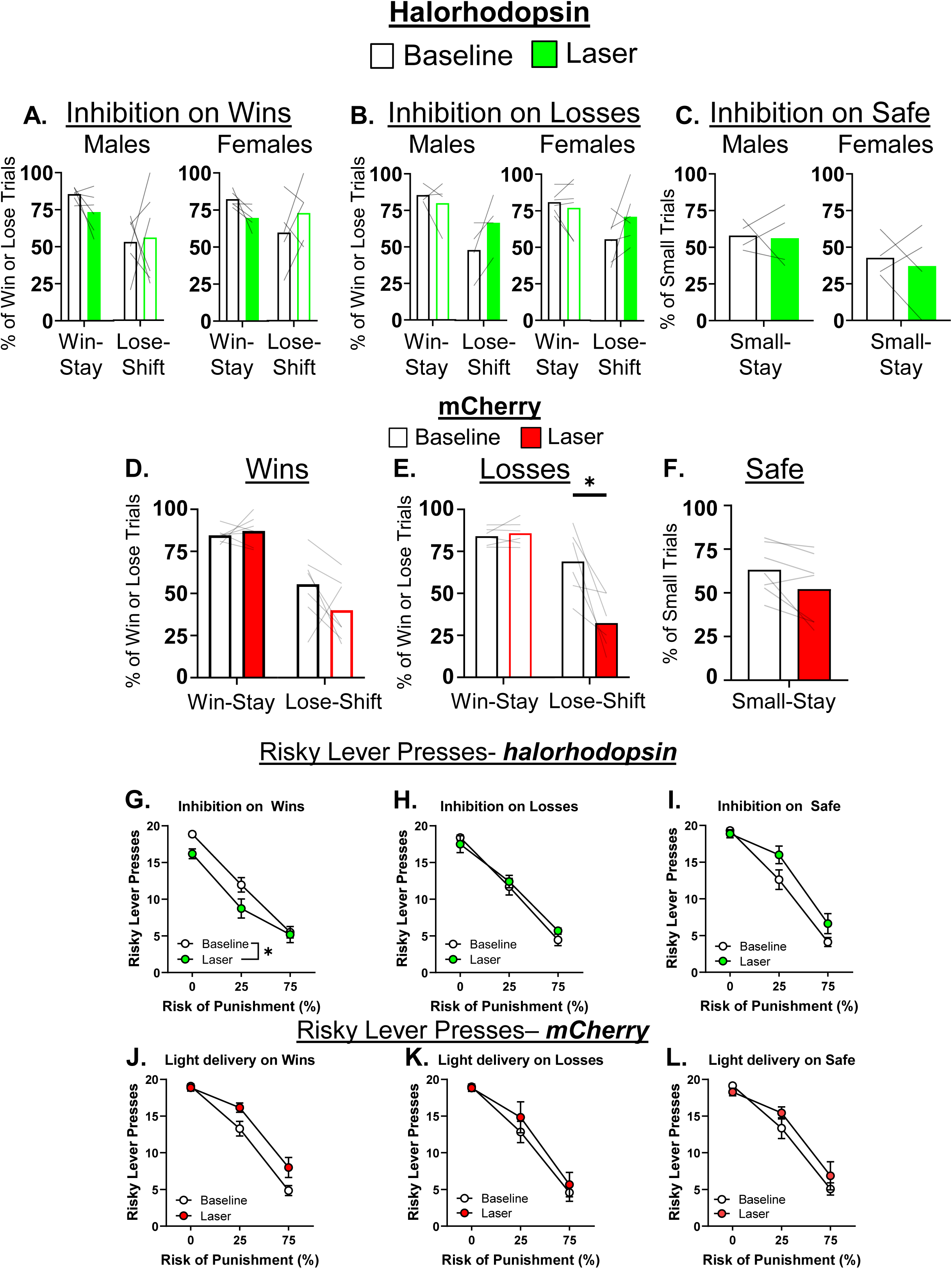
Supplemental figures for optogenetics experiments. **A)** Optogenetic results separated by sex for VTA dopamine neuron inactivation on Wins, **B)** Losses, and **C)** Safe outcomes. **D)** Comparisons of Win-Stay/Lose-Shift behavior in mCherry rats when laser was delivered on Win, **E)** Loss, and **F)** Safe outcomes. **G)** – **L)** Number of presses of the Risky lever when laser light was delivered on Win, Lose, and Safe outcomes in halorhodopsin (Halo) and mCherry rats. Grey lines on bar graphs indicate data from individual rats. * p < 0.05 for comparisons of light delivery condition.

